# Dynamics and variability of neuronal subtype responses during growth, degrowth, and regeneration

**DOI:** 10.1101/685917

**Authors:** Jamie A. Havrilak, Layla Al-Shaer, Noor Baban, Nesli Akinci, Michael J. Layden

## Abstract

**Background:** We are interested in nervous system dynamics in adult and regenerating animals. Preliminary studies suggest that some species alter neuronal number to scale with changes in body size. Similarly, in some species regenerates resulting from wholebody axis regeneration are smaller than their pre-amputated parent, but they maintain the correct proportionality, suggesting that tissue and neuronal scaling also occurs in regenerates. The cell dynamics and responses of neuronal subtypes during nervous system regeneration, scaling, and whole-body axis regeneration are not well understood in any system. The cnidarian sea anemone *Nematostella vectensis* is capable of wholebody axis regeneration, and its transparent, “simple” body plan and the availability of fluorescent reporter transgenic lines allow neuronal subtypes to be tracked *in vivo* in adult and regenerating animals. A number of observations suggest this anemone is able to alter its size in responses to changes in feeding. We utilized the *NvLWamide-like::mCherry* neuronal subtype transgenic reporter line to determine the *in vivo* response of neuronal subtypes during growth, degrowth, and regeneration.

**Results:** *Nematostella* alters its size in response to caloric intake, and the nervous system responds by altering neuronal number to scale as the animal changes in size. Neuronal numbers in both the endodermal and ectodermal nerve nets decreased as animals shrunk, increased as they grew, and the changes were reversible. Whole-body axis regeneration resulted in regenerates that were smaller than their pre-amputated size, and the regenerated nerve nets were reduced in neuronal number. Different neuronal subtypes had several distinct responses during regeneration that included consistent, no, and conditional increases in number. Conditional responses were regulated, in part, by the size of the remnant fragment and the position of the amputation site. Regenerates and adults with reduced nerve nets displayed normal behaviors, indicating that the nerve net retains functionality as it scales.

**Conclusion:** These data suggest that the *Nematostella* nerve net is dynamic, capable of scaling with changes in body size, and that neuronal subtypes display differential regenerative responses, which we propose may be linked to the scale state of the regenerating animals.

## Introduction

We are interested in understanding the dynamics of neuronal subtypes during whole-body axis regeneration and changes in body size. Typically, whole-body axis regeneration occurs through a combination of remodeling of existing tissue and growth of new tissue through cell proliferation. Regenerates resulting from whole-body axis regeneration are typically smaller than the pre-amputated animal but display correct proportionality [1–7], suggesting that rescaling of some tissues and organ systems is part of the normal regenerative response [4,7]. Many species are able to alter their adult size based on environmental conditions. For example, planarians grow and shrink in response to feeding and starvation, and the number of some neuronal subtypes in fixed planarians positively correlates with length [8–10]. This indicates that at least some nervous systems scale as adult body size is altered. However, tracking individual neuronal fates *in vivo* within the same individual in planarians is difficult. As such, investigating the dynamics of neuronal regenerative responses and scaling have been difficult to investigate.

The cnidarian sea anemone *Nematostella vectensis* possesses key features that make it well suited to investigate nervous system dynamics during growth, degrowth, and regeneration. *Nematostella* is diploblastic with a polyp bauplan typified by corals and anemones. The mouth, surrounded by tentacles, defines the oral end of the oral-aboral axis. It possesses a nerve net nervous system present in endodermal and ectodermal tissues that is easy to visualize using transgenic reporters because animals are optically clear at all life stages [2,11]. We previously identified the *NvLWamide-like::mCherry* transgene, which is expressed in five neuronal subtypes distributed along the oral-aboral axis in both the endoderm and ectoderm, within the pharynx, and in the tentacles (Figure 1A) [12,13]. During growth from juvenile to adult, at least two *NvLWamide-like* subtypes increase proportionally with length (Figure 1B,C) [13]. *Nematostella* alter their size in response to nutrient uptake. During periods of constant feeding they grow [14], and we noticed that animals left without food for a month or more were noticeably smaller, which is consistent with observations of the anemone *Actinia equina* that shrunk after being left unattended in labs for over two years during World War I [15]. However, the dynamics of *Nematostella* growth and degrowth have not been quantified.

**Figure 1:**
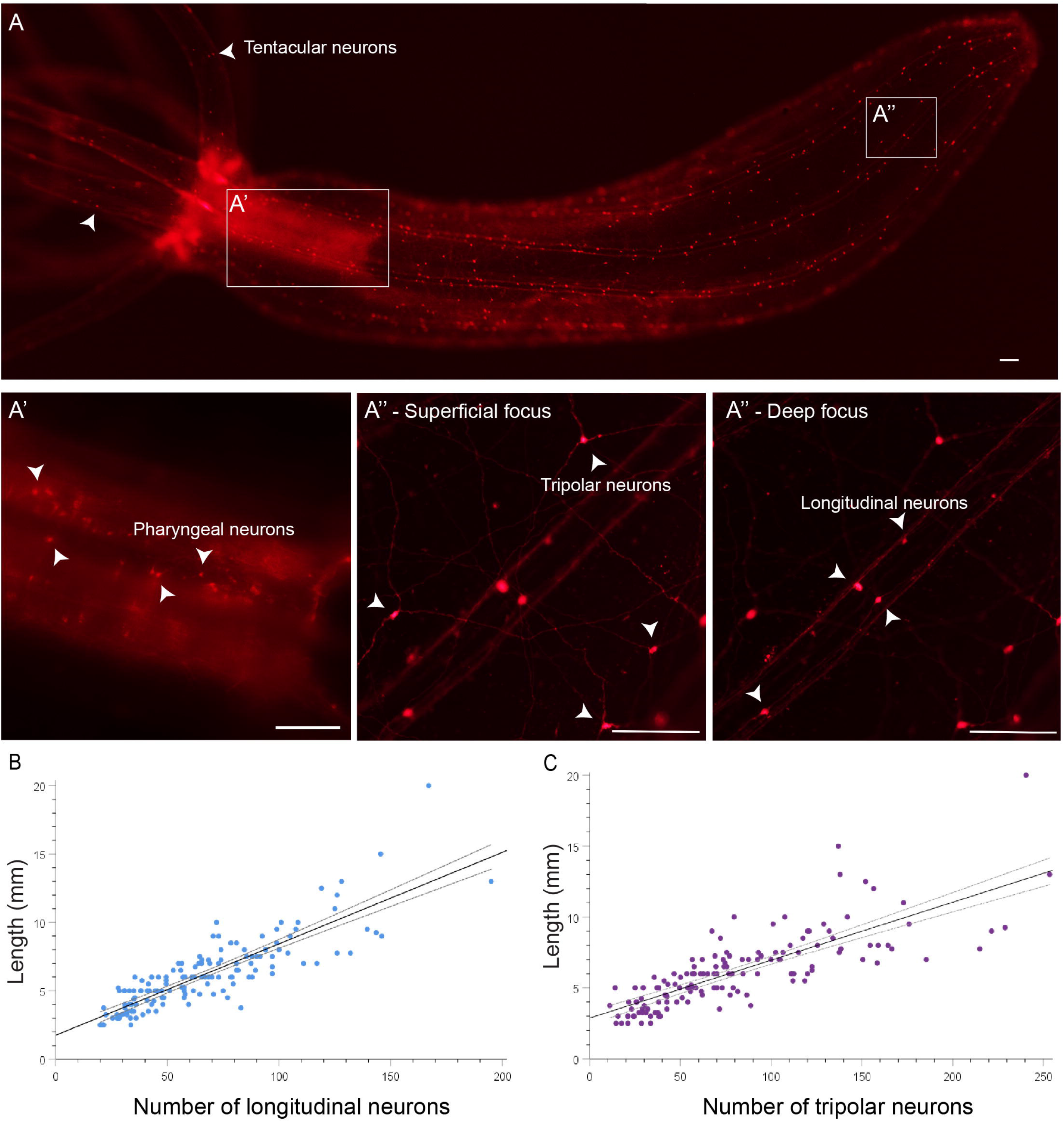
*NvLWamide-like::mCherry+* longitudinal and tripolar neuronal number positively correlate with length. (A) *NvLWamide-like::mCherry* positive animal showing the different neuronal subtypes used in this study. Arrowheads in A identify tentacular neurons. (A’) 10x magnification, showing pharyngeal neurons. (A’’) 20x magnification and different focal planes, demonstrating the location of neural subtypes present in the body column, within separate tissues. Tripolar neurons, located in the ectoderm, are captured in a superficial focal plane of the animal. Longitudinal neurons, residing in the endoderm, are captured in a deeper focal plane of the animal in the same location. Arrowheads identify neural subtypes labeled within each panel. (B-C) There is a strong positive correlation between animal length and the number of longitudinal (*r*_144_ = 0.854, *p* < 0.001, n = 146) and tripolar neurons (*r*_144_ = 0.791; *p* < 0.001, n = 145). Scale bars = 100μm.

Whole-body axis regeneration of the oral structures is well characterized in *Nematostella*. Oral structures regenerate in a stereotyped manner in 6-7 days [7,16–18]. Following amputation, a global increase in cell death enriched at the amputation site occurs from 1.5 to 12 hours after amputation [19]. The wound completes heals by 6 hours, and cell proliferation enriched at the amputation site is first detected at 12 hours post amputation. From 12 hours to 7 days post amputation cell proliferation in the blastema adds new endodermal and ectodermal tissue as the oral structures regenerate [7,17]. Regenerates are smaller than their pre-amputated adult, but like planarians and *Hydra* they are correctly proportioned [7,18,20]. The fact that proportionality is maintained despite the smaller size of regenerates suggests that regeneration combines both tissue regrowth and tissue remodeling during whole-body axis regeneration. Cell tracking confirmed that portion of the regenerated pharynx is derived from remodeled mesenteries, but the extent of tissue remodeling during *Nematostella* regeneration is not yet known [7]. The fact that *Nematostella* alters its size, regenerates are smaller than precut animals, and neurons within individuals can be tracked over time using transgenic animals makes *Nematostella* a potential model to determine how nervous systems respond during growth, degrowth, and whole-body axis regeneration.

Here we confirmed that *Nematostella* alters its size in response to feeding and tested the hypothesis that the nervous system scales with changes in size. Because regenerates are smaller than pre-amputated animals, we also tested whether or not the nerve net of regenerates is reduced and tracked the responses of individual neuronal subtypes during regeneration. To test these hypotheses, responses of *NvLWamide-like::mCherry+* neurons were quantified during body scaling throughout feeding, starvation, and whole-body axis regeneration. The *Nematostella* nervous system scales with changes in size by altering neuronal numbers and is reduced in regenerates. Despite differences in size, all nervous systems maintained known behaviors. Interestingly, we observed differential context dependent responses for one neuronal subtype during regeneration. Collectively, these data suggest that the *Nematostella* nerve net is highly dynamic and scales with body size. To the best of our knowledge these findings are the first direct confirmation that nervous systems adjust the number of neurons within individuals to maintain an appropriate scale state and are the first evidence that regenerating neuronal subtypes display variable responses during whole-body axis regeneration.

## Results

### *NvLWamide-like::mCherry+* neural subtypes used in this study

We assessed the potential for five previously described neuronal subtypes that express the *NvLWamide-like::mcherry* transgene (mesentery, tentacular, pharyngeal, longitudinal, and tripolar) to be used to track neuronal dynamics during nervous system scaling and/or regeneration in this study (Figure 1A) [13]. The ruffled structure of the mesenteries made reliable quantification of mesentery neurons almost impossible, and thus we excluded them from further characterization.

Pharyngeal and tentacular neurons can be scored for presence or absence in regenerating animals. Both neuronal subtypes are found within their namesake structures (Figure 1A,A’) [13]. In our previous study, accurate quantification of pharyngeal neurons was best achieved with fixation and confocal microscopy to overcome the autofluorescence of the pharynx (Figure 1A, A’) [13]. The need to fix animals prohibits tracking this neuronal subtype over time within live individuals. However, pharyngeal neurons are completely lost following amputation of oral structures, allowing their presence/absence to be scored during regeneration. We quantified the number of tentacular neurons in and across individuals. Tentacular neuronal number is not stereotyped between tentacles in a single individual or across individuals (Supplemental Tables 1 and 2). The lack of an obvious correlation between tentacular neuronal number and size, and the potential challenge of repeated identification of the same tentacle in individuals make them less than ideal for tracking dynamics with changes in animal size. However, similar to the pharyngeal neurons, all tentacles are lost following amputation and the presence or absence of tentacular neurons is scorable in regenerates.

Longitudinal and tripolar neurons are expressed along the oral-aboral axis and are well suited to track dynamics in generating or scaling nervous systems. Both of these subtypes increase in number as animals grow from juvenile to adult polyps (Figure 1B, C) and are easily quantified, because each neuronal subtype can be distinguished by their location and morphology (Figure 1A) [13]. Tripolar neurons have a columnar epithelial shaped soma, are located in the ectoderm, and have three neurites that project from the basal end of the cell originating at roughly equal intervals (Figure 1A”-Superficial focus) [13]. Longitudinal neurons are located in the endoderm within the eight paired longitudinal nerve tract bundles that run the length of the oral-aboral axis. In addition to being endodermal rather than ectodermal, they are easily distinguished from tripolar neurons because two neurites extend 180° to one another within longitudinal axon tracts and always along the oral aboral axis (Figure 1A” – Deep focus). Bisection of adult polyps results in remnant fragments that contain both longitudinal and tripolar neurons. Regeneration can be assessed by comparing the number of neurons present in remnant fragments to the neuronal number in regenerates. Thus, we can quantify dynamic changes of these neuronal subtypes during regeneration and in adults as they change size.

We conclude that four neuronal subtypes (tentacular, pharyngeal, longitudinal, and tripolar) allow us to assess regeneration and/or adult nervous system scaling.

### *NvLWamide-like+* longitudinal and tripolar neuron numbers correlate with polyp length

If tripolar and longitudinal neurons scale with length, their numbers should correlate with the length of adult polyps. To test this hypothesis, the length and neuronal numbers for 146 adult animals were plotted and the strength of association between these two factors was determined using Pearson’s correlation coefficients (Figure 1B,C). Both longitudinal and tripolar neurons positively correlated with length. However, the strength of the correlation between length and longitudinal neurons was slightly greater than that between length and tripolar neurons (*r*_144_ = 0.854, *p* < 0.001; *r*_143_ = 0.791; *p* < 0.001 respectively). We conclude that both longitudinal and tripolar neuron numbers correlate with animal length, and that these neuronal subtypes are suited to test the hypothesis that the *Nematostella* nerve net dynamically scales with changes in size.

### *Nematostella* undergoes growth and degrowth, and neuronal number scales with changes in size

To determine if at least portions of the *Nematostella* nerve net scale with changes in body size, animal length and neuronal number were followed in individuals undergoing growth and degrowth. We used two feeding regimes: starved and then fed (starved-then-fed) or fed and then starved (fed-then-starved). The initial design was to track length and neuronal numbers in replicate experiments over fourteen weeks. Animals would be either starved or fed for seven weeks, then the feeding regimen would be switched for the remaining seven weeks. However, after three weeks of feeding, many of the fed-then-starved animals became large and accurately counting neurons within the time constraints of our paralytic would be challenging if animals continued to grow. We therefore switched the animals to starvation for the remaining 11 weeks (Figure 2; Supplemental Figure 1; black lines). To account for variable lengths of animals at the start of the experiment, all changes in length were normalized to percent length for reporting purposes.

**Figure 2:**
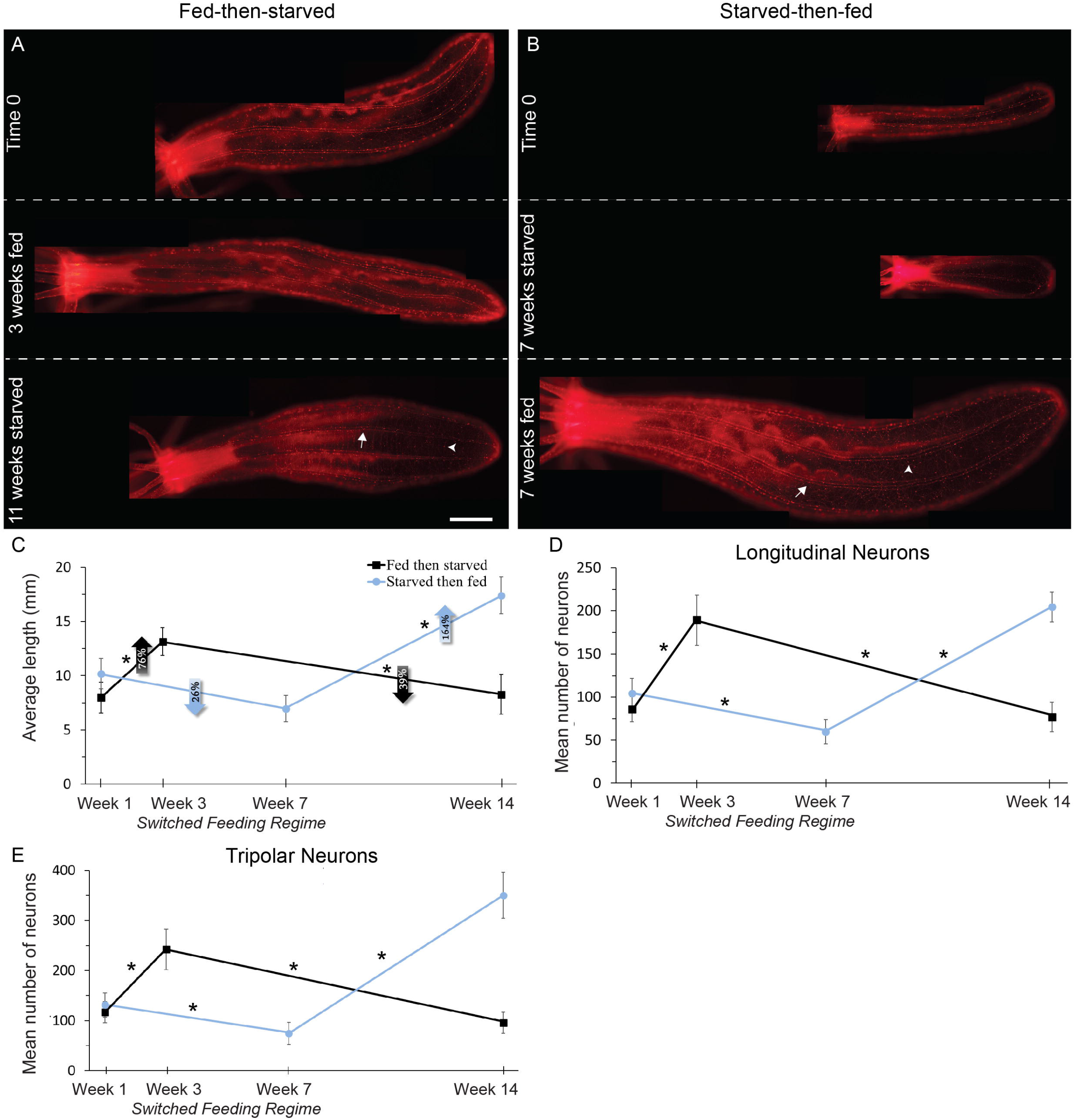
The *Nematostella* neuronal subtypes scale with changes in size. (A-B) Visualization of *NvLWamide-like* neurons in relation to body size during periods of feeding and starvation. Panels in A and B are each of a single individual. Arrows indicate longitudinal neurons and arrowheads indicate tripolar neurons. Oral end is to the left. Scale bar = 1mm. (C) Increases and decreases animal length due to feeding and starvation, respectively. (D) Longitudinal neurons increase and decrease during periods of feeling and starvation, respectively. (E) Tripolar neurons increase and decrease following periods of feeding and starvation, respectively. (C-E) See Supplemental Table 5 for full model results (C: *F*_2,34_ = 25.75,*p* < 0.001, *η*_p_^2^ = 0.60; D: *F*_2,36_ = 77.35,*p* < 0.001, *η*_p_^2^ = 0.81; E: Tripolar, *F*_2,36_ = 91.91, *p* < 0.001, *η*_p_^2^ = 0.84). Data points are means ± SEM, n = 10 animals per treatment, \**p* ≤ 0.05. Scale bar = 1mm.

Relative to their starting length, the fed-then-starved animals grew significantly by 75.9% ± 20.8% after three weeks (Figure 2A,C; black line; *p* = 0.008). Relative to their length at the time of their switch to starvation, the fed-then-starved animals significantly decreased in length by 38.6% ± 9.1% after 11 weeks (Figure 2A,C; black line; *p* = 0.008). In the fed-then-starved group, the longitudinal and tripolar neurons significantly increased (Longitudinal: *p* < 0.001; Tripolar: *p* < 0.001) and decreased (Longitudinal: *p* < 0.001; Tripolar: *p* < 0.001) in number respectively as animals grew and shrank in length (Figure 2D,E; black lines).

Relative to their starting length, the starved-then-fed animals decreased in length by 26.1% ± 7.4% after 7 weeks of starvation, which was trending towards significance (Figure 2B,C; blue line, *p* = 0.098). Relative to their length when switched to feeding, the starved-then-fed group significantly increased in length by 164%± 34.9% (Figure 2B,C; blue line; *p* < 0.001). Similar to the previous group, the number of longitudinal and tripolar neurons significantly increased *(p* ≤ 0.001 for both) and decreased (Longitudinal: *p* = 0.007; Tripolar: *p* = 0.026) as animals grew and shrank, respectively (Figure 2 D,E; blue lines).

Resting *Nematostella* polyps are elongated with tentacles extended. However, they are rarely observed at their maximum extension, and whether their resting length is invariant has not been quantified. To be confident that we were observing growth and degrowth rather than variable states of resting elongation, we repeatedly measured the length of single animals across the size spectrum daily for 5 continuous days. Regardless of animal size the resting length did vary, and the average variability was 14% ± 3% over 5 days (Supplemental Figure 2A). As expected, the number of longitudinal neurons did not change over the same 5-day period (Supplemental Figure 2B; *p* = 0.145).

The observed decreases in length of 26.1% and 38.6% over seven and eleven weeks, respectively, and increases in length of 75.9% and 164% for three and seven weeks, respectively (Figure 2C) are outside of the 14% error rate for measuring length at a single time point. We conclude from these data that *Nematostella* increases and decreases length depending on caloric intake, and that the nerve net responds by altering neuronal numbers to scale with size.

### Starved animals maintain characteristic behaviors

To determine if starved animals retain normal behaviors with a reduced nerve net, we repeated our starvation experiments and tested the polyp’s ability to perform known behaviors before and after starvation (Figure 3). As predicted, longitudinal and tripolar numbers were significantly decreased following starvation (Figure 3A,B; *p* < 0.001 for both). Known behaviors for *Nematostella* are: they produce peristaltic waves by generating contractions along the oral-aboral axis; they capture and ingest prey; they retract in response to mechanical stimuli; and they produce gametes in response to light and temperature cues [14,21,22]. Animals starved for seven weeks or eleven weeks performed peristaltic waves, albeit at slightly reduced rates (Figure 3C,D; starved seven weeks: *p* = 0.005; starved 11 weeks: *p* = 0.014). To test their ability to retract from a mechanical stimulus, we developed a poking assay to test an animal’s ability to retract when poked at its oral end (see Methods). Prior to starvation animals responded to touch 89.5% and 100% of the time. After 7 and 11 weeks of starvation animals responded 89.5% and 83.3% of the time, respectively (Figure 3E-J; Table 1; Supplemental Movies 1 and 2). Reproductive ability was also tested in animals starved for 7 weeks. 6/6 females and 5/6 males produced and released gametes both before and after starvation. These findings argue that the reduced nerve net retains functions across scale states.

**Figure 3:**
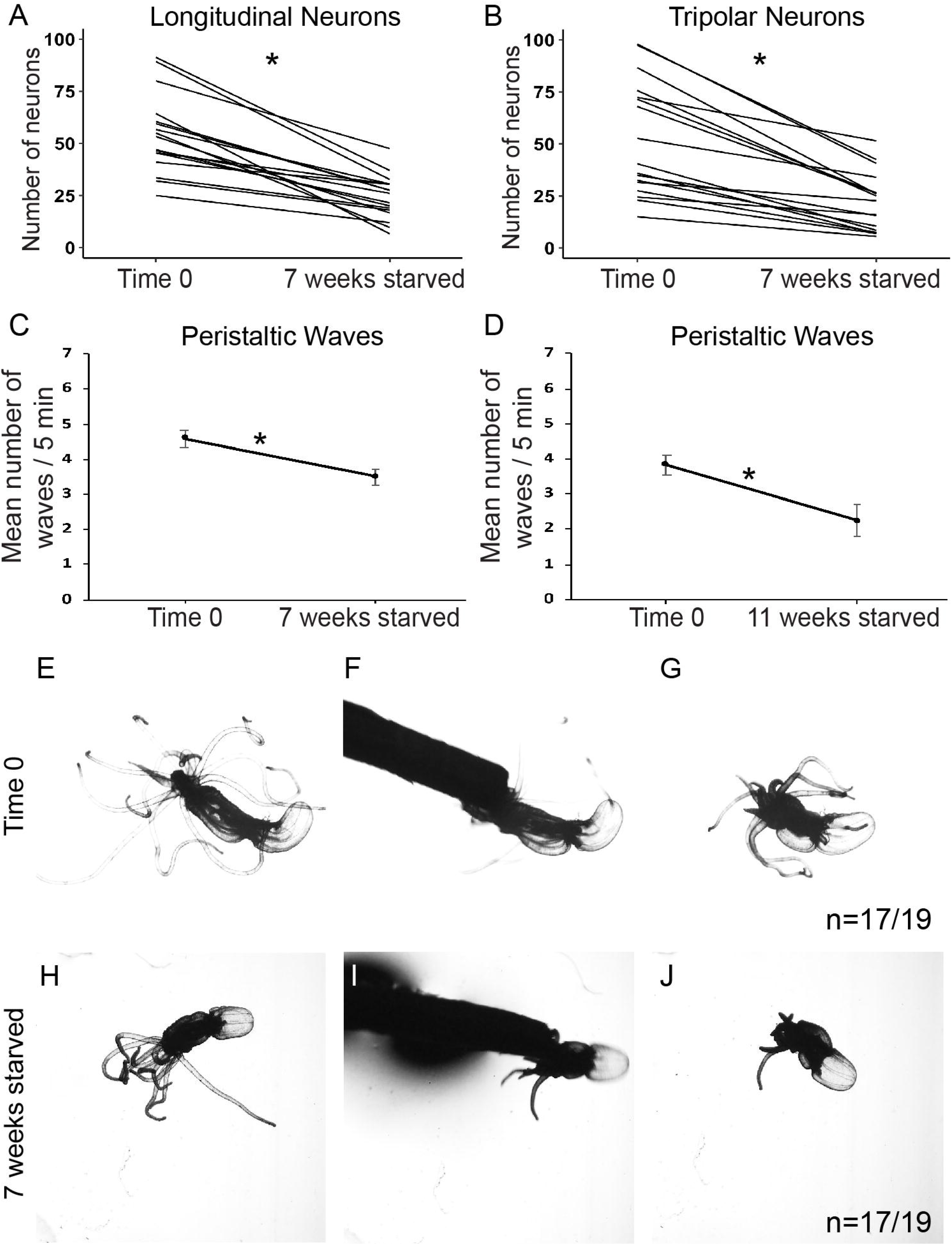
Starved animals maintain characteristic behaviors. (A-B) Quantification of the number of longitudinal and tripolar neurons in individual animals following each poking experiment. The numbers of both longitudinal and tripolar neurons significantly decreased after 7 weeks starvation (I: *t*_16_ = 7.97, *p* < 0.001; J: *t*_16_ = 6.99, *p* < 0.001; Both n = 17). (C-D) Peristaltic waves still occurred in all animals starved for 7 and 11 weeks (C: *t*_11_= 3.45, *p* = 0.005; D: *t*_11_= 2.92, *p* = 0.014; Both n = 12). (E-G) A normally fed animal responded to the touch stimulus from a toothpick by retracting its tentacles and oral region. (H-J) Following 7 weeks starvation, the same animal again responded to being touched with a toothpick by retracting its tentacles and oral region. Data points are means ± SEM, \**p* ≤ 0.05.

**Table 1:** Summary of the analysis for the action-reaction poking assay in starved and regenerated animals.

| Experimental treatment | Timepoint |  | Number to respond to touch stimulus | Percent to respond to touch stimulus |
| --- | --- | --- | --- | --- |
| Starvation | Before: 7 weeks starved |  | 17/19 | 89% |
|  | After: 7 weeks starved |  | 17/19 | 89% |
|  | Before: 11 weeks starved |  | 12/12 | 100% |
|  | After: 11 weeks starved |  | 10/12 | 83.3% |
| Regeneration | Uncut | Small | 5/5 | 100% |
|  |  | Medium | 6/6 | 100% |
|  |  | Medium-large | 5/5 | 100% |
|  |  | Large | 6/6 | 100% |
|  | 7 dpa | Small | 5/5 | 100% |
|  |  | Medium | 6/6 | 100% |
|  |  | Medium-large | 5/5 | 100% |
|  |  | Large | 6/6 | 100% |
|  | 14 dpa | Small | 5/5 | 100% |
|  |  | Medium | 6/6 | 100% |
|  |  | Medium-large | 5/5 | 100% |
|  |  | Large | 6/6 | 100% |

### Tentacular and pharyngeal neurons regenerate

To investigate regeneration, we wanted to bisect animals into equal *oral* and *aboral* remnants (Figure 4A, see Methods). To ensure animals were equally bisected we developed a method to quantify the amputation site using longitudinal neurons. Longitudinal, but not tripolar, neurons are evenly distributed amongst equal quadrants along the oral-aboral axis (Supplemental Figures 2C,D,3F; Longitudinal: *p* = 0.33; Tripolar: *p* < 0.001). Thus, comparing the number of longitudinal neurons present in pre- and post-cut animals immediately after bisection ensured amputation occurred at the intended position (Supplemental Figure 4A-D, compare uncut timepoints with time 0 cut).

**Figure 4:**
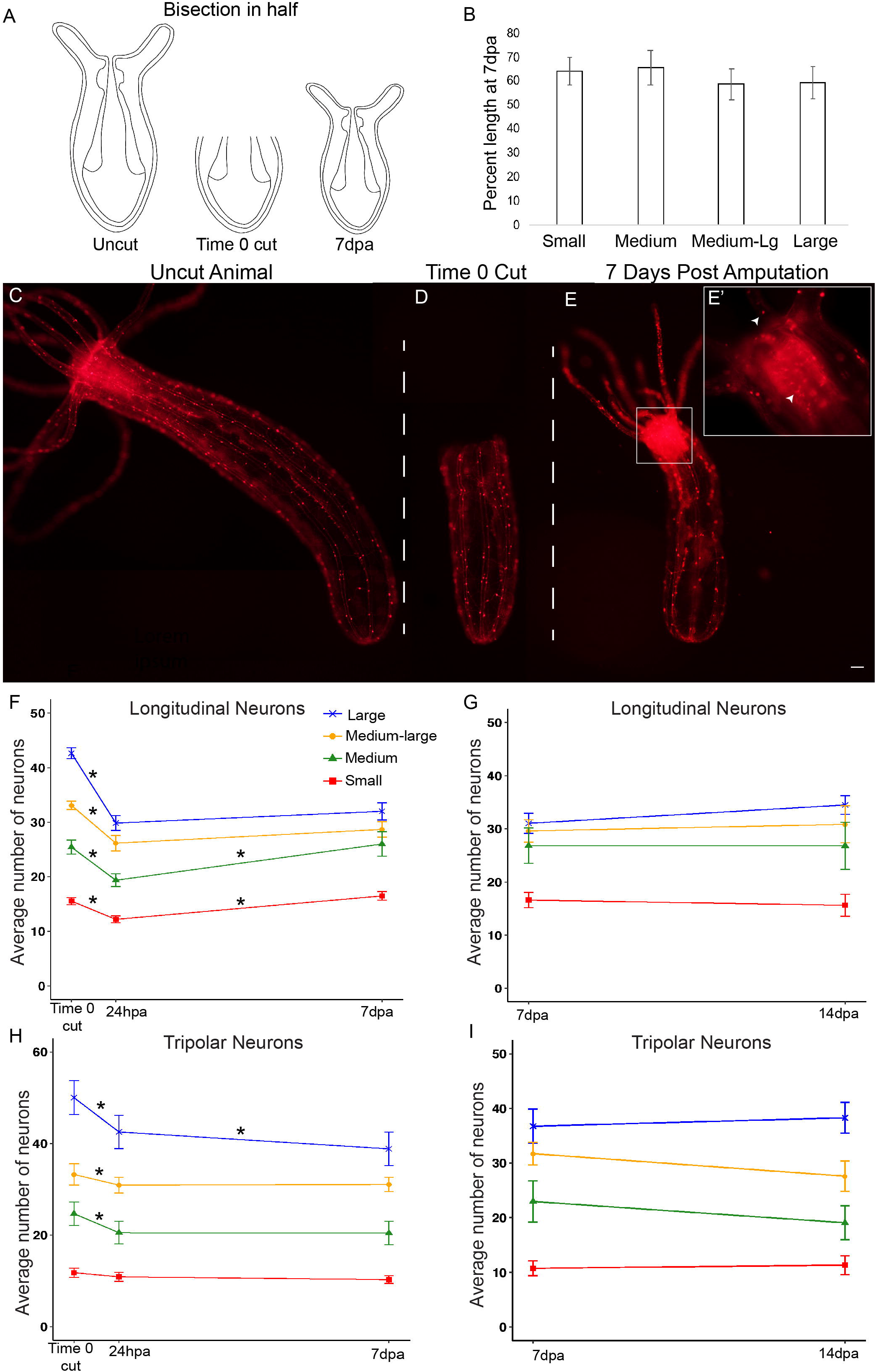
*NvLWamide-like* neuronal subtypes have differential responses during regeneration. (A) Schematic for oral bisection at ~50% body length. (B) Percent length regained in 7 day post amputation (dpa) animals bisected in half did not differ between animals of different size categories (*F*_3,27_ = 0.23, *p* = 0.875, *η*_p_^2^ = 0.025, n = 11 small; 8 medium; 6 medium-large; 6 large). (C) Uncut *NvLWamide-like::mCherry* expressing animal. (D) Same animal as C at time 0 cut. (E) Same animal as C and D at 7dpa, with regenerated oral structures. Scale bar = 100μm. (E’) 10x magnification of the boxed oral region in E, top and bottom arrowheads indicate tentacular and pharyngeal neurons respectively. (F-I) Quantification of longitudinal and tripolar neurons in regenerating animals. Lines represent the average number of neurons per radial segment in small (red), medium (green), medium-large (orange), and large (blue) animals. (F) Longitudinal neurons regenerated back to numbers similar to time 0 cut in the small *(p* = 0.99) and medium sized animals *(p* = 1.00), but did not regenerate in the larger animals by 7dpa (medium-large: *p* = 0.003, large: *p* < 0.001; *F*_5.1,134.4_ = 13.46,*p* < 0.001, *η*_p_^2^ = 0.34; n = 29 small, 20 medium, 16 medium-large, 18 large). (G) Allowing animals to regenerate another 7 days did not increase longitudinal neuron number in animals of any size category (*F*_3,42_ = 0.86, *p* = 0.47, *η*_p_^2^ = 0.06; n = 12 small; 12 medium; 10 medium-large; 12 large). (H) Tripolar neurons did not regenerate by 7dpa (*F*_4.5,116.2_ = 3.93,*p* = 0.003, *η*_p_^2^ = 0.13; n = 28 small, 19 medium, 16 medium-large, 18 large). (I) Allowing animals to regenerate another 7 days did not increase tripolar neuron number in animals of any size category (*F*_3,42_ = 2.04, *p* = 0.122, *η*_p_^2^ = 0.13). Regardless of size, there was a near significant decrease in tripolar neurons between 7dpa and 14dpa (*F*_1,42_ = 3.97, *p* = 0.053, *η*_p_^2^ = 0.09; n = 12 small; 12 medium; 9 medium-large; 11 large). See Supplemental Table 5 for full model results. Data points and bars represent means ± SEM, \**p* ≤ 0.05.

Consistent with previous studies, oral structures, including the mouth, pharynx, and tentacles, regenerated in the *aboral* remnant by 7dpa in 82/83 individuals and regenerates were smaller than their pre-amputated parent (Figure 4B-E; Supplemental Figures 3A-E,4E) [7,17,18,23]. We also quantified the length of regenerates at 7 days post amputation (dpa) and compared it to their pre-cut length. Based on unexpected size dependent responses of longitudinal neurons during regeneration (described below), we grouped animals based on starting size for the remainder of our analyses. Animals were categorized as small (20-40 longitudinal neurons and ~2.5-5 mm), medium (41-60 longitudinal neurons and ~3.5-5.5 mm), medium-large (61-80 longitudinal neurons and ~5-7.5mm), and large (81-100 longitudinal neurons and ~6.5-8.5 mm). At 7dpa regenerates from small and medium animals bisected in half were slightly longer, relative to the length of the precut animal, than regenerates from medium-large and large animals, but this difference was not statistically significant (Figure 4B; Supplemental Figure 4E; *p* = 0.875).

We next assessed the responses of individual neuronal subtypes. Pharyngeal and tentacular neurons were typically observed in their namesake structures by 72 hours post amputation (hpa) and were identified in 100% of regenerated animals at 7dpa (Figure 4E; Supplemental Figure 3A-E). We conclude that both tentacular and pharyngeal neurons regenerated by 7dpa, and that their regeneration coincided with the re-appearance of their corresponding tissues.

### Longitudinal and tripolar neurons have variable responses in regenerating animals

To determine if longitudinal and tripolar neurons regenerated by 7dpa, we compared the number of neurons at 7dpa in *aboral* remnants immediately following amputation (time 0 cut) and at 24hpa, which is after the wave of cell death [19] (Figure 4; Supplemental Figure 4). As mentioned above, all regenerates were initially treated as one group. However, we noticed a size dependent variability in responses observed for longitudinal neurons during regeneration, which motivated us to track how neuronal numbers changed during regeneration of different size groups.

Regardless of size, there was a decrease in longitudinal neuron number from time 0 cut to 24hpa (Figure 4F, Supplemental Figure 4A,B; *p* < 0.001 for all size groups). The number of longitudinal neurons significantly increased between 24hpa and 7dpa in small and medium animals (Figure 1F, red and green lines; *p* < 0.001 for both). The number of longitudinal neurons in 7dpa regenerates was statistically similar to the numbers observed in time 0 cut animals for both small and medium groups (small: *p* = 0.99, medium: *p* = 1.00). In contrast, longitudinal neurons did not increase from 24hpa to 7dpa in medium-large and large animals between 24hpa and 7dpa (Figure 1F, yellow and blue lines; *p* = 0.25 and *p* = 0.34, respectively). To address the concern that larger animals had not completely regenerated by 7dpa, we performed additional replicates and allowed animals to regenerate to 14dpa. The additional time did not increase longitudinal numbers from those observed at 7dpa for any size category (Figure 4G; *p* = 0.47 for all). We conclude that regenerates have less longitudinal neurons than their pre-amputated parents, which is consistent with their reduced size (Figure 4B, Supplemental Figure 4E). Additionally, we observed a size dependent response during regeneration where an increase in longitudinal number from 24hpa to 7dpa occurred in small and medium sized animals, but not larger animals.

In contrast to longitudinal neurons, tripolar neurons did not increase in number in any group (Figure 4H; Supplemental Figure 4C,D). In small animals, tripolar neuron number was indistinguishable at time 0 cut, 24hpa, and 7dpa (Figure 4H, red line; *p* = 1.00 for all comparisons), indicating that small regenerates did not lose tripolar neurons by 24hpa, nor did they gain any by 7dpa. In medium, medium-large, and large animals, the number of tripolar neurons decreased between time 0 cut and 24hpa (Figure 4H; *p* = 0.001, *p* = 0.013, and*p* < 0.001, respectively). Medium and medium-large animals did not increase the number of tripolar numbers between 24hpa and 7dpa *(p* = 1.00 for both), and large animals further decreased tripolar neurons by 7dpa *(p* = 0.055). Allowing regenerates to continue to 14dpa did not increase tripolar neuron number compared to 7dpa counts (Figure 4I; *p* = 0.12).

We conclude that during the initial regeneration of oral structures in bisected *Nematostella*, there are at least three distinct neuronal responses. The first is neuronal regeneration that occurs 100% of the time (e.g. pharyngeal and tentacular neurons), and the second is neurons that do not respond or regenerate (e.g. tripolar neurons). The third is a variable response, where neurons are decreased in number over the first 24hpa but only return to the number present immediately post-amputation in a subset of animals (e.g. longitudinal neurons increase in number between 24hpa and 7dpa to numbers similar to the time 0 cut fragment in small and medium animals).

### Amputation site and size of *aboral* remnant impacts the regeneration of longitudinal neurons

Possible explanations for the variation in longitudinal neuron responses are that either the size of the remnant fragment or the size of the pre-amputated animal influenced their variable responses during regeneration. To test these hypotheses, the amputation sites of animals of different sizes were shifted to generate larger and smaller remnant fragments. If the size of the remnant fragment influences regeneration, the shifted amputation site should alter the regenerative response. However, if pre-amputation size influences the initial regenerative response, then altering the amputation site should not impact the longitudinal neuron response.

Smaller *aboral* fragments were generated from large animals by shifting the amputation site aborally to remove 75% of the oral-aboral axis (Figure 5A). The resulting remnant fragment resembled the remnant of a medium animal bisected in half (Figure 5C,D, compare black and faded green lines), and interestingly the number of longitudinal neurons increased between 24hpa and 7dpa (Figure 2C, *p* = 0.046). Thus, altering the cut site of large animals to generate smaller remnant fragments shifted the longitudinal neurons to now respond and increase in number from 24hpa to 7dpa.

**Figure 5:**
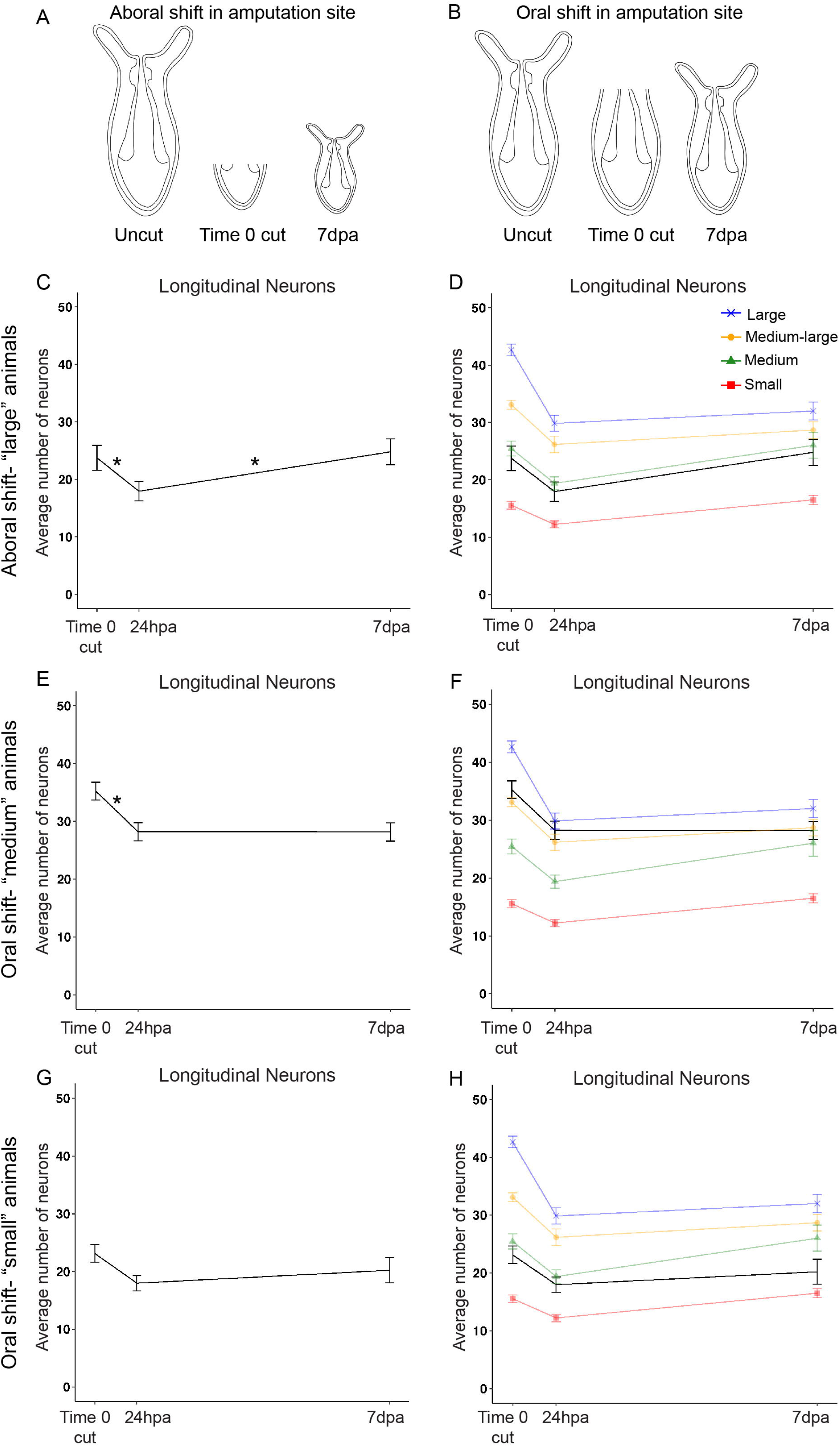
Differential regenerative responses of longitudinal neurons are partially dependent on the size of the remnant fragment. (A-B) Schematics demonstrating the shifts in amputation sites used to obtain remnant fragments of varying sizes. (C-H) Quantification of longitudinal neurons in regenerates with shifted amputation sites at time 0 cut, 24hpa, and 7dpa. (C) Regenerates from an aborally-shifted amputation site in large animals regenerated their longitudinal neurons *(p* = 0.046; *F*_2,14_ = 5.58, *p* = 0.017, *η*_p_^2^ = 0.44, n = 8), resembling regenerates of small and medium sized animals bisected at the midline of the oral-aboral axis (D). (E) Regenerates from an orally-shifted amputation site in medium animals did not regenerate their longitudinal neurons *(p* = 1.00; *F*_2,24_ = 21.67, *p* < 0.001, *η*_p_^2^ = 0.64, n = 13), resembling regenerates from large animals bisected at the midline of the oral-aboral axis (F). (G) Regenerates from an orally-shifted amputation site in small animals did not regenerate their longitudinal neurons *(p* = 0.47; *F*_2,24_ = 8.56, *p* = 0.002, *η*_p_^2^ = 0.42, n = 13), which is different from medium animals bisected at the midline of the O-A axis (H). See Supplemental Table 5 for full model results. Data points in C-H represent means ± SEM, \**p* ≤ 0.05.

Orally shifting the amputation site of medium sized animals to remove only 25% of the oral-aboral axis (Figure 5B) generated remnant fragments similar to those generated by bisecting medium-large or large animals into equal halves (Figure 5E,F, compare black and faded blue and yellow lines). In this case, no increase in the number of longitudinal neurons was observed between 24hpa and 7dpa *(p* = 1.00).

When the amputation site of small animals was orally shifted (Figure 5B), the *aboral* remnant resembled a medium animal bisected into equal halves (Figure 5G,H, black lines). While the average number of longitudinal neurons did increase between 24hpa and 7dpa, the change was not statistically significant (Figure 5G,H, compare black and faded green lines; *p* = 0.47). It should be noted that neuronal numbers increased in some but not all regenerates. The variability in longitudinal neuron increase in small animals with orally shifted amputation sites was distinct from other experiments where the entire population more or less responded similarly (Supplemental Figures 4B, 5A-C). Regardless, these data suggest that the pre-amputation size of the animal was not a factor regulating the responses observed for longitudinal neurons during regeneration. Remnant fragment size did influence longitudinal neuron responses. However, longitudinal neurons in small animals with orally shifted cut sites did not respond, suggesting additional factors influence longitudinal neuron responses.

### Form and function are restored in regenerated *Nematostella* by 7 days post amputation

We next sought to determine if normal functions are also restored by 7dpa, which would argue that the nervous system is functional following the initial regenerative response. At 7dpa and 14dpa, animals were able to catch and digest prey (Figure 6A-C; Movies 3 and 4). Regenerates observed at 7dpa also had 4-5 peristaltic waves on average over a 5-minute interval (Figure 6A-C, arrows; Movie 5), with no statistical difference in the number of waves between regenerates of different size categories (Figure 6D; *p* = 0.35). This number is similar to the number of waves observed in uncut animals (t-test: *t*11 = 1.10, *p* = 0.30), and in previously published observations [21]. We next examined poking-induced behaviors in pre- and post-regenerated animals of all size categories. In all cases, the same animal responded to the poking assay pre-bisection, 7dpa, and 14dpa 100% of the time (Figure 6E-L; Table 1; Movies 6 and 7). It should be noted that the function of longitudinal and tripolar neurons remains unknown, and thus we could not directly test behaviors regulated by those neurons. Nevertheless, both form and function were restored by 7dpa, which suggests that regenerates have completed whole-body axis regeneration.

**Figure 6:**
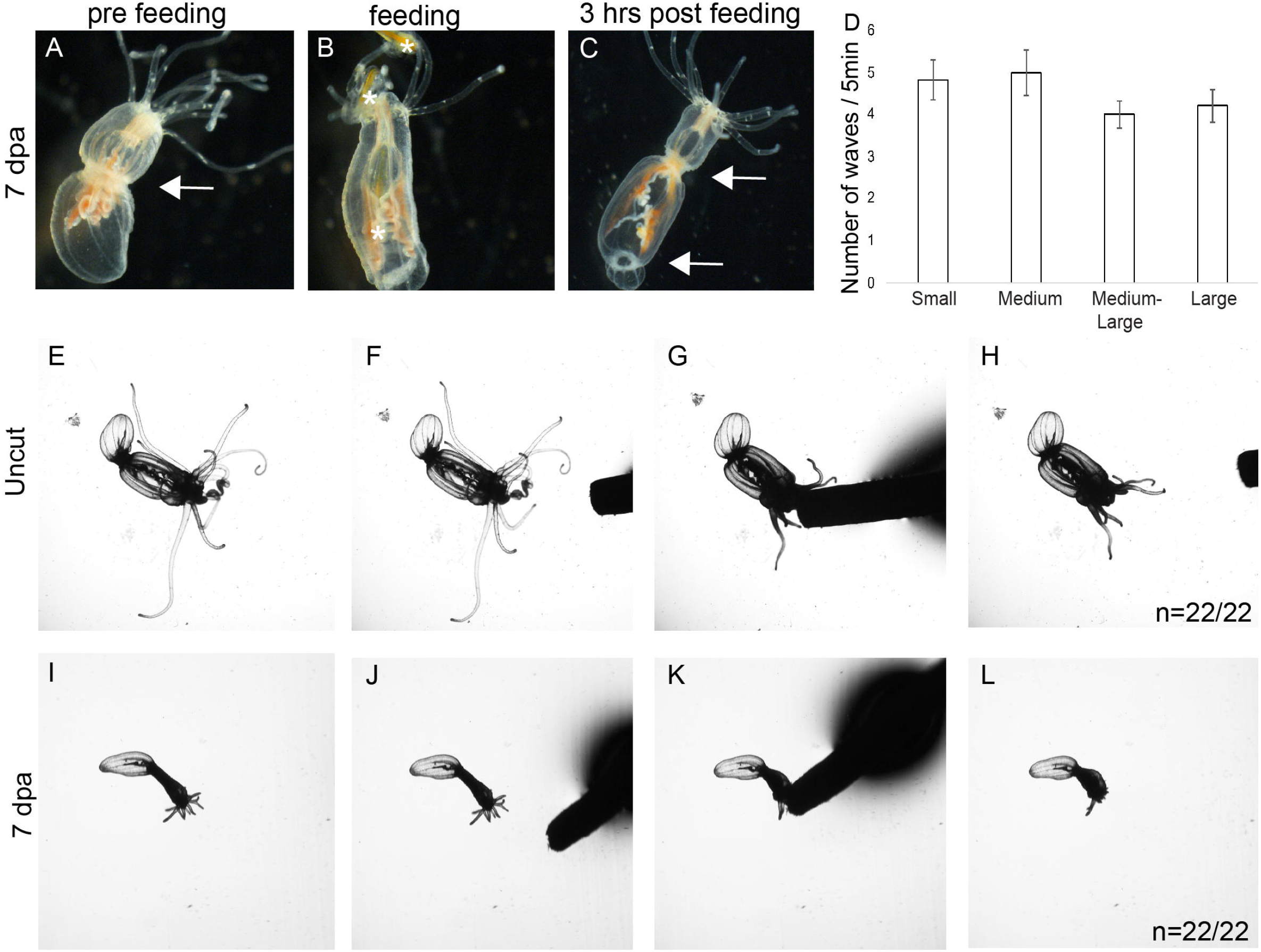
Normal behaviors are restored in regenerated animals by 7 days post amputation (dpa). Representative regenerated animal pre-feeding (A), during feeding (B), and 3 hours post feeding at 7dpa (C). Asterisks identify the locations of brine shrimp during capture, ingestion, and digestion. Note the increase in orange pigment in the mesenteries due to the digestion of the brine shrimp in C. All regenerates were capable of capturing, ingesting, and digesting prey (n =6/6 small; 5/5 medium; 5/5 medium-large; 6/6 large). Arrows in A and C highlight peristaltic wave movements. (D) Animals in all size categories carried out similar numbers of peristaltic waves by 7dpa (*F*_3,17_ = 1.16, *p* = 0.35, *η*_p_^2^ = 0.17). (E-L) Response to the poking assay is maintained in regenerated animals (n = 5/5 small, 6/6 medium, 5/5 medium-large, 6/6 large). (E-H) *Nematostella* responded to the touch stimulus by retracting its tentacles when touched with a toothpick. (I-L) At 7dpa, the same animal again responded to being poked with a toothpick by retracting its tentacles.

## Discussion

### *Nematostella* alter size and neuronal number in response to caloric intake

We demonstrated that *Nematostella* decrease and increases in length in response to starvation and feeding, respectively. This is not surprising, as most *Nematostella* researchers have been aware of this phenomenon from anecdotal observations, planarians are known to undergo growth and degrowth during feeding and starvation, and anemones left unattended in a laboratory during World War I survived without food or water changes, but were dramatically reduced in size [8,9,15,24–27]. In addition to the data presented here, a subpopulation of animals left unattended in our lab for well over three months during COVID-19 restrictions were dramatically reduced in size, but survived and regrew once fed. Our data confirmed that *Nematostella* is able to alter the size of the adult polyp, making this a viable system to investigate scaling of adult animals.

Neurons within the endoderm and ectoderm modulate their number to positively scale with length. Animals gain or lose neurons as they increase and decrease in length, respectively. Importantly, changes in neuronal number are reversible, which argues that the nervous system senses and responds to changes in size to maintain the proper proportion of neuronal cells. Neuronal scaling in planarians has been implied by the observation that the number of visual and brain neurons correlate with body length in fixed animals [8]. Recent reports in *Hydra,* a cnidarian relative of *Nematostella,* suggest that neuronal scaling may occur in a number of cnidarian species [28]. Interestingly, both *Hydra* and planarians are capable of whole-body axis regeneration, and their regenerates are also smaller than the precut animal, suggesting that neuronal scaling is a common phenomenon in animals whose adult size is dynamic.

Our data suggest that reduced nervous systems retain normal function. Polyps retract in response to being poked, catch and digest prey, coordinate peristaltic waves, and are able to perform reproductive behaviors. We infer from our studies that, regardless of scale state, the animals are able to perform complex behaviors. For example, both starved and fed animals are capable of distinguishing between a potential predator (toothpick) and prey (brine shrimp), because while the animals responded to the poking assay with a defensive retraction, they readily captured and ingested food. Despite the limited number of functional assays in *Nematostella,* the range of behaviors that are unaffected by the total number of neurons suggests that the nervous system is functional in all scale states. Similarly, we are unaware of any evidence that smaller planarians have behavioral deficits compared to larger animals. Collectively these observations indicate that neuronal scaling without changing behaviors occurs in both cnidarians and bilaterian species.

The most likely mechanism for altering neuronal number is through regulation of cell proliferation and cell death. Planarians alter their total cell numbers during growth and shrinking by modulating gene expression to regulate the balance between cell death and proliferation [25–27,29]. Fed *Nematostella* show signs of increased proliferation [30], and in other cnidarians increases in cell proliferation are associated with growth and tissue expansion [17,30–32]. Alternatively, cell death in response to nutrient deprivation is well documented in animals. While future efforts should confirm that regulating proliferation and cell death are critical components of scaling, the interesting questions are: how is neuronal number monitored; and what mechanisms regulate programs to increase and decrease neuronal number as animals change in size?

### Nervous system scale state may influence the regeneration of neuronal subtypes

The fact that nervous systems scale with size in *Nematostella,* and that regenerates are smaller than pre-cut animals, suggests that the differential regenerative responses of individual neurons may be linked to the scale state of the regenerate. The two neuronal subtypes that we confirmed have reduced numbers in regenerates and scale with size are tripolar and longitudinal neurons. These two subtypes had variable responses during regeneration (no regeneration and a conditional increase in neuronal number, respectively). This suggests that the “acceptable range” of neurons needed to maintain the correct scale state varies for different subtypes. Support for this is that the variability of tripolar neurons was much higher than longitudinal neurons for each size group, and the distribution of tripolar neurons along the oral-aboral axis was not as uniform as longitudinal neurons, suggesting that there is more flexibility in the number and distribution of tripolar neurons required for proper function than exists for longitudinal neurons.

One criticism is that the variable responses reflect incomplete regeneration. However, this is unlikely in this study for multiple reasons. First, form and function were both restored in animals by 7dpa. Although the nervous system was reduced in overall number, all behaviors were present. More importantly, when nervous systems were reduced during starvation in the feed/starve experiments, their functions were not lost, suggesting that behaviors were maintained in nervous systems regardless of scale state. Second, no increase in neuronal number was observed when animals were allowed to regenerate for an additional week (14dpa). By 14dpa, animals had not been fed for almost three weeks. In the feed/starve experiment, the first noticeable decrease in neural number during starvation occurred in week three. Thus, we suspect that extended regeneration without eating would not improve neuronal regeneration. We do expect that feeding regenerates would increase both tripolar and longitudinal neurons. However, we would not characterize this as regeneration because the same phenomenon is observed in fed un-lesioned animals. Third, our feed-starve data and evidence from other anemones suggest that adult sea anemones, including *Nematostella,* do not have a fixed size, but rather alter their size based on caloric intake [28,31,33]. Thus, form and function can be restored in regenerates that are smaller than their precut parent, and regeneration does not require returning to a precut length. This seems to be a common feature of animals capable of whole-body axis regeneration, as many of them also have adults whose size is dynamic. The fact that 7dpa animals possessed the ability to feed, digest food, and display other known behaviors strongly indicates that regeneration had completed. We are confident that *Nematostella* possesses neurons that display differential responses during regeneration of oral structures.

### Differential regeneration of neuronal subtypes during whole-body axis regeneration

The conditional response of longitudinal neurons is interesting. The data suggest that smaller remnants need to generate new neurons for the regenerated nervous system to be at the correct scale state, whereas larger remnants do not. Current explanations for why are speculative, but the mechanisms of regeneration may vary based on the size of the remnant fragment and amputation site. Unlike *Hydra, Nematostella* regeneration requires cell proliferation [7,17]. However, evidence suggests that remodeling of remnant tissue also occurs [7]. EdU labeling indicates that larger animals still utilize cell proliferation during regeneration (Supplemental Figure 6), but perhaps they shift the balance between remodeling and proliferation to favor remodeling. This idea is supported by our observation that, on average, regenerates generated from small and medium animals bisected in half tended to be slightly longer than regenerates from medium-large and large animals when normalized to their pre-amputated length. An alternative hypothesis is that the scale state relationship of longitudinal neurons to length is not linear across all sizes, with larger animals adding new neurons at a slower rate. In this case, the larger remnant fragments have a sufficient number of neurons to be within the acceptable range for that scale state. To distinguish between these models will require identifying a paralytic that also causes *Nematostella* to stretch to their maximal extended length. This would minimize potential variability in length measurements and allow researchers to 1) improve the resolution and accuracy of the ratios for length to neuron number, and 2) determine if smaller remnants regenerate more new tissue than longer remnants.

The failure of longitudinal neurons to increase in number in small animals with orally shifted amputation sites challenges both of these models, but it does not eliminate them. The remnant fragment was small enough that an increase longitudinal number was predicted by the “remnant fragment size influences regenerative response” model. Intriguing to us was the fact that the responses of longitudinal neurons in small animals with orally shifted amputation sites was much more heterogeneous than the other experiments (Supplemental Figures 4,5). The orally shifted amputation sites in small animals were proximal to the mouth and pharynx, whereas other amputations performed in this study were much more distal to the oral structures. It is possible that oral amputations close enough to the mouth resemble loss of a particular structure (i.e. amputation of limb) rather than whole-body axis regeneration, and subsequently the regenerative mechanisms are distinct. This could explain the variability in our data, and it offers additional hypotheses to test in the future.

## Conclusion

The ability to observe and quantify neuronal subtypes makes *Nematostella* a powerful model to investigate neurogenesis in the context of both regeneration and whole-body scaling. Our data suggest that the regenerative responses of neuronal subtypes in *Nematostella* are variable during regeneration of oral structures, and that the nervous system scales with size. We suspect that variable neuronal responses are linked to regenerates rescaling their nervous system to accommodate their smaller size, but additional studies are necessary to validate this hypothesis. How *Nematostella* control the regeneration and incorporation of new neurons into the existing nerve net, and how they scale the numbers of these neurons to body size in real time remain unknown, but both are exciting areas for future investigation. Lastly, our data imply that experiments aimed at uncovering the molecular mechanisms regulating neural regeneration in *Nematostella* require careful consideration of the differential responses of individual neuronal subtypes during regeneration when designing experiments and interpreting results.

## Methods

### Animal care

All animals were kept at room temperature and fed rotifers or crushed brine shrimp until they reached the young juvenile adult stage, at which time they were transferred to 17°C in the dark and fed brine shrimp 4x/week, unless otherwise noted for experimental purposes. *Nematostella* were maintained in 1/3x artificial sea water, *(Nematostella* medium) at a pH of 8.1-8.2, that was changed once per week.

*Nematostella* were relaxed in 7.14% (wt/vol) MgCl2 in *Nematostella* medium for ~10 minutes prior to length measurement, neural quantification, imaging, or bisection. No paralytic was used during behavioral experiments. The lengths of relaxed animals were measured in mm with a ruler placed under a Petri dish containing the animal.

### Quantification and imaging of *NvLWamide-like::mCherry*-expressing animals

To quantify longitudinal and tripolar neuron numbers, we first determined that the number of neurons in each radial segment were similar to one another within each individual animal (Supplemental Tables 3 and 4). The minimal variation allowed us to quantify neuronal number in an individual by calculating the representative number of longitudinal and tripolar neurons present by averaging 2-4 segments for each animal/time point (Supplemental Figure 3F). The same individual was kept isolated so that the number of neurons present within one animal could be tracked over time.

*NvLWamide-like::mCherry* tripolar and longitudinal neurons were counted on live animals under a dissection microscope (Nikon SMZ1270), on live animals placed under a slide with a raised coverslip on a compound microscope (Nikon NTi), or on full-length images of *Nematostella* captured on a Nikon Nti. All quantifications were done by hand, because it was more reliable than images acquired on a fluorescent compound scope and these animals were too large to generate confocal montages covering the whole animal in 20 minutes or less, which is the upper limit of the amount of time these animals tolerate the paralytic with no detrimental effects.

### Imaging

Live images of *NvLWamide-like::mCherry*-expressing animals were taken on a Nikon NTi with a Nikon DS-Ri2 color camera with Nikon Elements software. Large fulllength images are composed of several images stitched together. Confocal images were captured using a Zeiss LSM 880 with LSM software and processed using Imaris 8.4.1 software (Bitplane LLC) to create 3D images from serial optical sections (z-stacks). Images were then cropped and assembled using Photoshop and Illustrator (Adobe).

Live video was captured on a Nikon Eclipse E1000 microscope with Nikon Imaging software or a Nikon dissection microscope (Nikon SMZ1270) with a USB camera attachment. Movies were processed with ImageJ/Fiji software and screen shots were obtained from individual frames, then cropped and assembled using Adobe Photoshop and Illustrator.

### Regeneration experiments

For the quantification of longitudinal and tripolar *NvLWamide-like::mCherry* expressing neuronal subtypes during regeneration, only animals containing 20-100 longitudinal neurons were used, which were typically 2.5 to 9 mm long (Supplemental Figure 4A,B,E). This size range represents the sizes of naturally occurring wild-caught animals found along the East Coast of the United States (Adam Reitzel, University of North Carolina, Charlottesville, NC, personal communication, 11/2018).

Longitudinal and tripolar neurons were quantified first on the uncut animal, then immediately following bisection (time 0 cut), 24 hours post amputation (hpa), and 7 days post amputation (dpa). For some regenerating animals, additional counts were conducted at 14dpa. Animals were bisected in half with the amputation site at the oral-aboral midpoint (~50% body removed), bisected below the pharynx (“oral shift”, ~25% removed), or bisected at the aboral end (“aboral shift”, ~75% removed).

We developed a strategy to ensure the amputation sites were reproducibly positioned in the middle 50% of the animal, oral 75% of the animal, or aboral 25% of the animal. Longitudinal neurons are equally distributed along the oral-aboral axis (Supplemental Figure 2C). Thus, confirmation of the cut sites were achieved by comparing the number of neurons in the remnant fragment immediately after amputation with the number of total neurons present in the precut animal.

Following quantification and/or bisection, the remnant *aboral* fragment was washed with *Nematostella* medium, then placed in the dark at room temperature. Prior to the start of the experiment, animals were starved for 3-4 days [7,17]. Animals were not fed for the duration of the regeneration experiments.

### Feed/Starve experiments

For each animal, the number of longitudinal and tripolar neurons were counted at the beginning of the experiment, at the time of their feeding regime switch (Starved-then-fed: 7 weeks later; Fed-then-starved: 3 weeks later), and at the end of the 14 week experiment (Starved-then-fed: 7 weeks later; Fed-then-starved: 11 weeks later). All animals were maintained on the same normal feeding schedule until the start of the experiment. At this point, animals were randomly assigned to either the starved-then-feed group or the fed-then-starved group, and their initial starting neural counts were taken. Animals on the feeding regime were fed brine shrimp 2x/week plus oyster 1x/week. All animals were given weekly water changes.

### Poking assay

Animals were transferred to individual dishes containing *Nematostella* medium and observed under a dissection microscope. Once the animal had relaxed its tentacles, a toothpick was slowly inserted into the medium so as to not disturb the animal prematurely. The animal was then “poked” with the tip of the toothpick at the oral region. The entire assay was video recorded and analyzed to determine the animal’s response. Following the “poking assay”, the animal was left alone in the dish until it relaxed again, upon which time the poking assay was performed again. The assay was performed 3 consecutive times. A positive response was determined if the animal responded to being touched with a toothpick by retracting some or all of its tentacles, scrunching or retracting in its mouth (oral region), or retracting its entire body. To be considered responsive, the animal had to react in at least 1 out of the 3 poking assays performed. The poking assay was done in a repeated fashion, with each animal being tested at the beginning and end of the experimental treatment.

### Peristaltic Waves

Animals were individually placed in dishes filled with *Nematostella* medium and observed under a dissection scope. Data collection did not begin until animals relaxed their tentacles and/or at least 75% of their body-column. Once relaxed, animals were observed for 5 continuous minutes and the number of peristaltic waves that occurred during this time was recorded. A peristaltic wave was defined as an inward radial contraction that propagated down the body column in an oral to aboral direction [21]. Waves that were in progress at the start of data collection were counted, and so were waves that did not go to completion by the end of the observation period. In the regeneration experiment, the number of waves observed/5 minutes was compared between animals categorized as small, medium, medium-large, and large. In the feed/starve experiment, animals were observed at the beginning and end of either a 7 or 11-week starvation period.

### Feeding behavior

Animals were transferred to individual dishes filled with *Nematostella* medium and observed using a dissection microscope. Once an animal relaxed its tentacles, freshly hatched brine shrimp (5-10) were added to the dish. Animals were first observed for their ability to capture and ingest brine shrimp. Once at least one was captured, the rest of the brine shrimp were removed and the animal was left overnight. The following day, the dish was checked for evidence of expelled brine shrimp that had not been digested following capture and ingestion.

### Reproductive behavior

Animals were kept individually in dishes filled with *Nematostella* medium. Each individual was spawned prior to the starvation period to determine their gametic sex. Those that released eggs were designated as female and those that released sperm were designated as male. Egg clutches are visible to the naked eye and are therefore easily scored for their presence or absence. Sperm release was determined by observing if a clutch of eggs from an isolated known female would be fertilized (scored by the presence of cleaving embryos) when given water from an unidentified individual’s dish. As a false-positive control, half of each known female’s eggs were set aside and observed to make sure no development occurred. Experimental animals were starved for 7 weeks while control animals were fed freshly hatched brine shrimp once per week over 7 weeks. All animals were kept in the dark at 17°C and received weekly water changes. After 7 weeks, all animals were placed on a light box in order to stimulate reproductive behavior. Whether or not animals released gametes was observed using the methods described above.

### EdU Stain

EdU incorporation was carried out as previously described [17]. Animals were incubated with 330uM EdU in *Nematostella* medium for 30 minutes, then fixed as previously described and dehydrated to 100% MeOH and stored at −20°C. Following rehydration, samples were permeabilized in PBT (PBS with 0.5% Triton X-100, Sigma) for ~2 hours. Animals were then treated with the reaction cocktail (prepared following the Click-It protocol provided by the manufacturer, with the exception of doubling the amount of Alexa Fluor azide) at room temperature in the dark for 20 minutes. Following 5 long washes in PBT, the animals were labeled with propidium iodide for 1 hour while rocking, washed 5X in PBT, and finally transferred to 90% glycerol for imaging.

### Statistical analysis

Two replicate data sets were initially collected for the feed-starve experiment by two different researchers. These data sets were combined for analysis because there was no significant difference in how individuals responded across all time points based on the researcher that collected the data (mixed ANOVA – time point x researcher: Length: *F*_2,34_ = 0.08, *p* = 0.92; Longitudinal: *F*_2,36_ = 0.52, *p* = 0.60; Tripolar: *F*_2,36_ = 0.60, *p* = 0.46). For the regeneration and feed-starve experiments, mixed analysis of variance (ANOVA) tests were used to analyze each dependent variable (i.e. body length, and longitudinal and tripolar neurons). For both experiments, each observation of an individual over time served as the within-subject factor (Regeneration experiment: time 0 cut, 24hpa, 7dpa; Feed/starve experiment: week 1, week of feeding regime switch, week 14). The between subject factors used were animal size category (small: 20-40mm, medium: 41-60mm, medium-large: 61-80mm, large: 81-100mm) and assigned feeding group (Feed-Starve or Starve-Feed) for the regeneration and feed/starve experiments, respectively. To determine if regeneration of longitudinal and tripolar neurons was complete by 7dpa, additional mixed ANOVAs were used to compare neuron counts at 7dpa and 14dpa by size category. A repeated measure ANOVA was used to determine if longitudinal and tripolar neurons were evenly distributed within equal body column quadrants. Data used for ANOVA tests met assumptions of normality and homogeneity. Greenhouse-Geisser corrected F-statistics are reported if sphericity was violated (i.e. Figure 4F,H). When omnibus ANOVA results were significant, post-hoc testing was performed using Bonferroni corrections in order to maintain tight control over the type 1 error rate. Effect sizes are reported as partial eta^2^ (*η*_p_^2^). Full ANOVA model results are reported in Supplemental Table 5 for all significant omnibus analyses. Paired t-tests were used to compare individual changes in the number of peristaltic waves, longitudinal and tripolar neurons, or length at two time points. All data were analyzed using SPSS version 25.0.

## Acknowledgements

We would like to thank Aldine Amiel and Eric Röttinger for critically reading this manuscript and providing feedback, as well as their personal communications regarding cell death in starved animals. We would like to thank Amber Rice and Bridget Dever for their input on the appropriate statistical methods to evaluate our data. We also thank Adam Reitzel for providing insights about what size ranges we should target to represent adult animals found in the wild. We thank David Balli and Omar Ahmed for providing help with R software and construction of the line graphs. This work was funded by the National Institutes of Health Award Number R01GM127615.

---

**Supplemental Figure 1:**
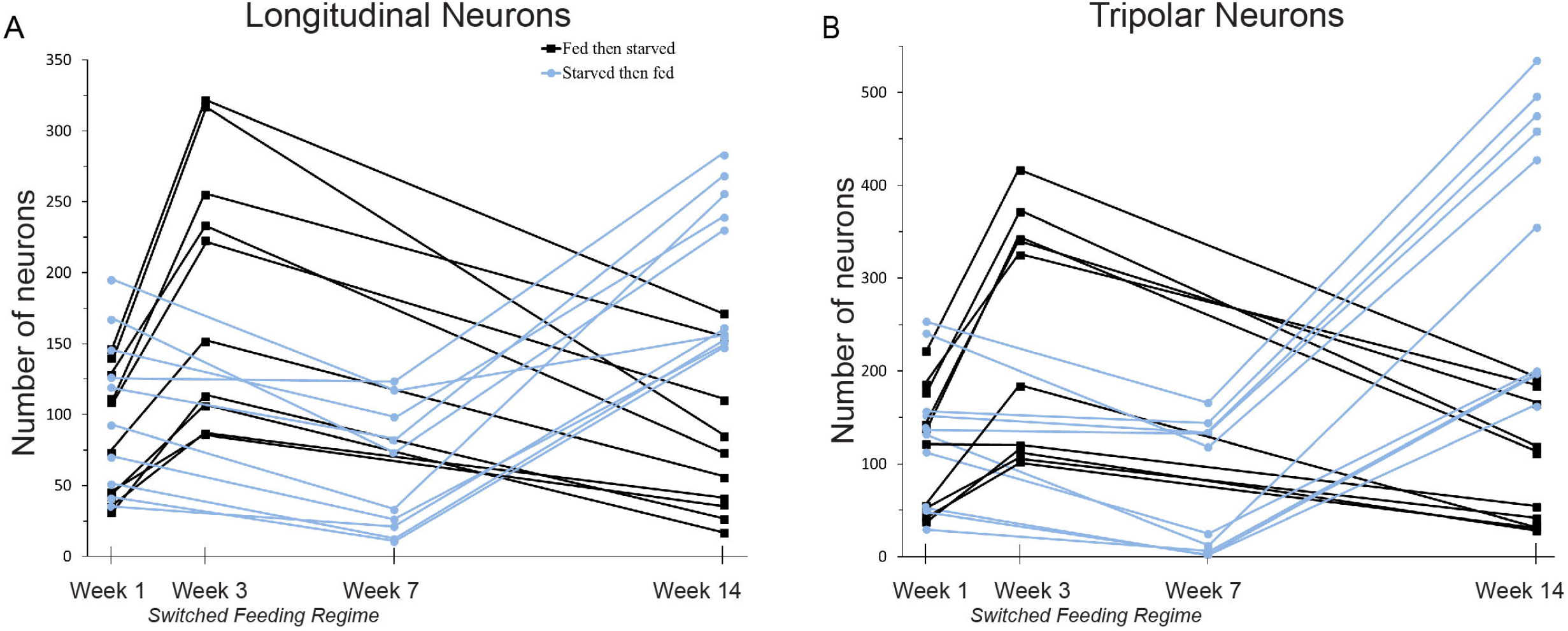
Individual longitudinal and tripolar neuron data from the feed/starve experiment. (A-B) Neurons were quantified in twenty individuals (10/treatment) at the start, time of feeding regime switch, and end of the feed/starve experiment. These data were used to determine the average responses of fed-then-starved and starved-then-fed animals, which are shown in Figure 2. See Supplemental Table 5 and Figure 2 for statistical analyses.

**Supplemental Figure 2:**
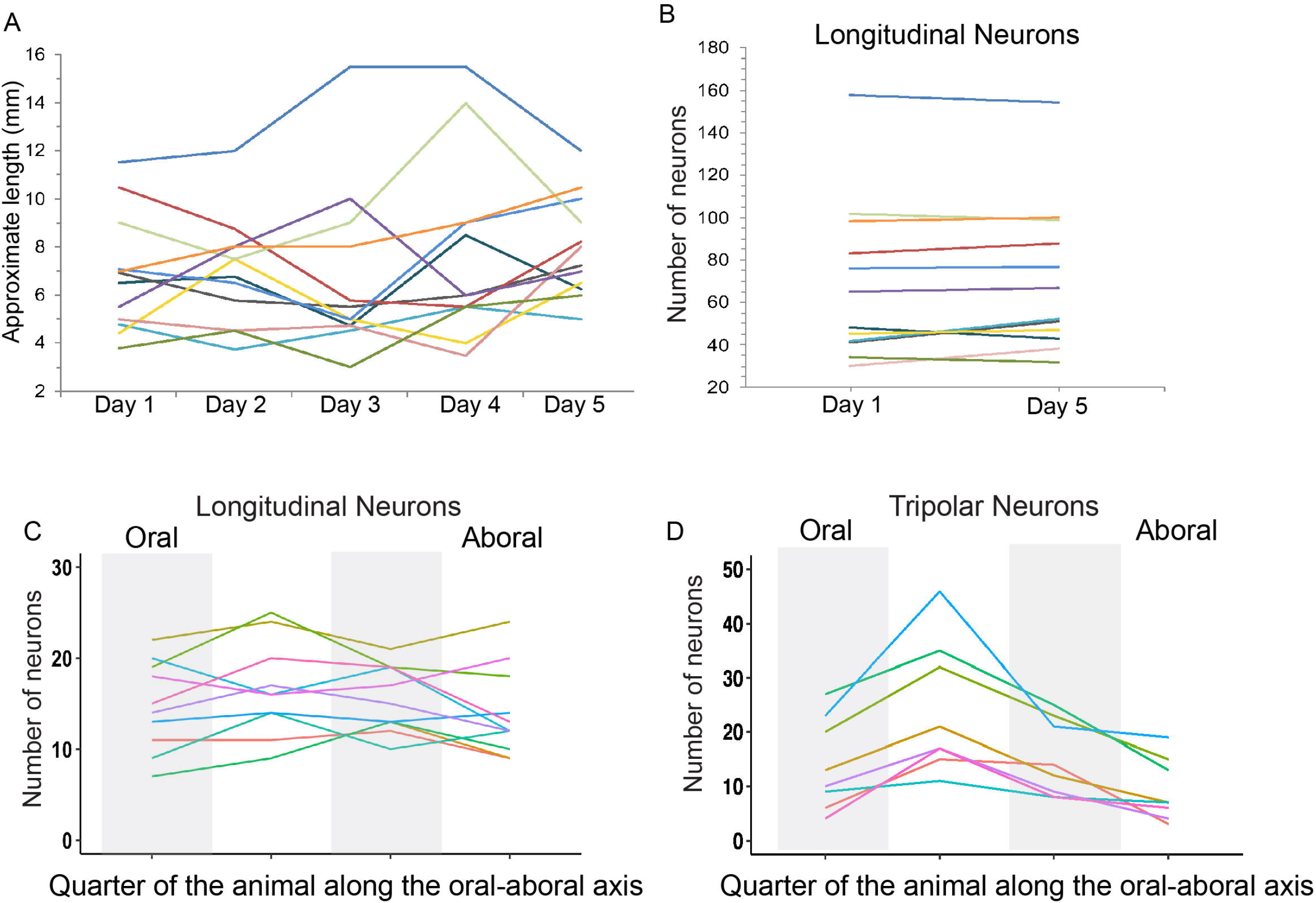
Length measurements and neural quantifications in *Nematostella*. (A) The average variability in the measured length of the same animal over 5 consecutive days was 14% ± 3%. (B) The number of longitudinal neurons did not change over the same 5 day period in the same animals as A (*t*_11_ = −1.57, *p* = 0.145). (CD) Quantification of the number of longitudinal and tripolar neurons along the oral-aboral axis in each of the 4 equal quarters. Gray and white bars distinguish the 4 quadrants. (C) Longitudinal neurons are equally distributed along the oral-aboral axis (*F*_3,45_ = 1.18, *p* = 0.33, *η*_p_^2^ = 0.73; n = 10), but tripolar neurons are not (*F*_3,24_ = 20.97, *p* < 0.001, *η*_p_^2^ = 0.72; n = 8). Each line represents an individual animal.

**Supplemental Figure 3:**
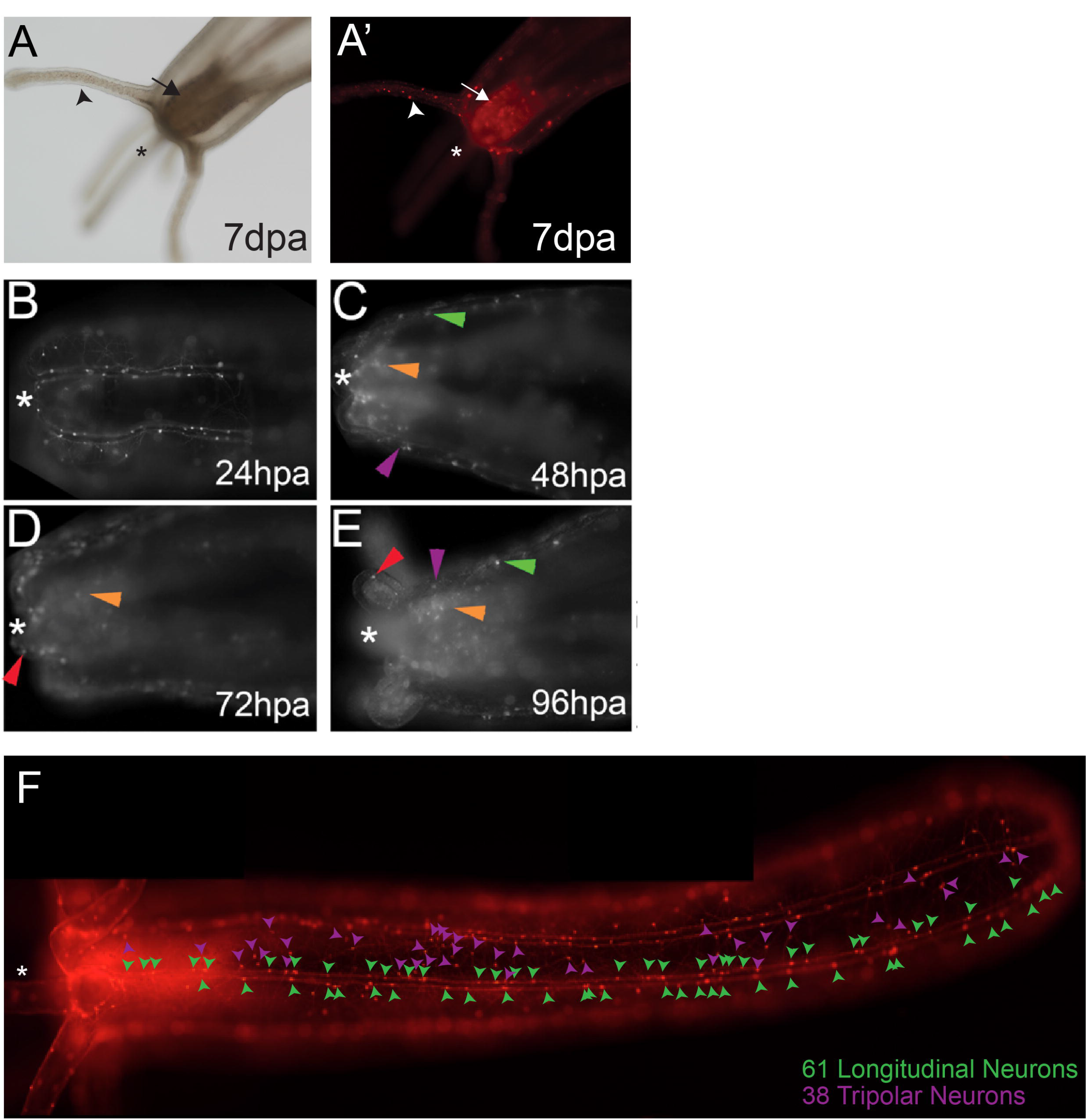
Regeneration of oral structures and neurons in *NvLWamide-like::mCherry* animals following bisection along the oral-aboral axis. (A) Regenerated oral structures, including a mouth, pharynx (arrow), and tentacles (arrowhead) at 7dpa. (A’) New neurons are observed in the regenerated oral structures, including pharyngeal (arrow) and tentacular (arrowhead) neurons. (B) Remnant fragment at 24hpa. No new structures were observed at this time point. (C) Tentacle buds were visible in the remnant fragment at 48hpa. Longitudinal (green arrowhead) and tripolar (purple arrowhead) neurons were observed. Regenerated pharyngeal neurons were also present in the regenerating pharynx (orange arrowhead). (D) Pharyngeal (orange arrowhead) and tentacular (red arrowhead) neurons were clearly visible in these regenerating structures by 72hpa. (E) Clearly regenerated pharyngeal (orange arrowhead) and tentacular (red arrowhead) neurons are seen at 96hpa. Longitudinal neurons (green arrowhead) and tripolar neurons (purple arrowhead) also populated the regenerated tissue, but whether these were regenerated neurons or from the remnant was undetermined. (F) Example of how longitudinal (green arrowheads) and tripolar (purple arrowheads) were quantified within a longitudinal track and radial segment, respectively. Note that for regeneration and feed-starve experiments, neuron counts were performed live under a dissection scope and not from images. Asterisks indicate oral opening.

**Supplemental Figure 4:**
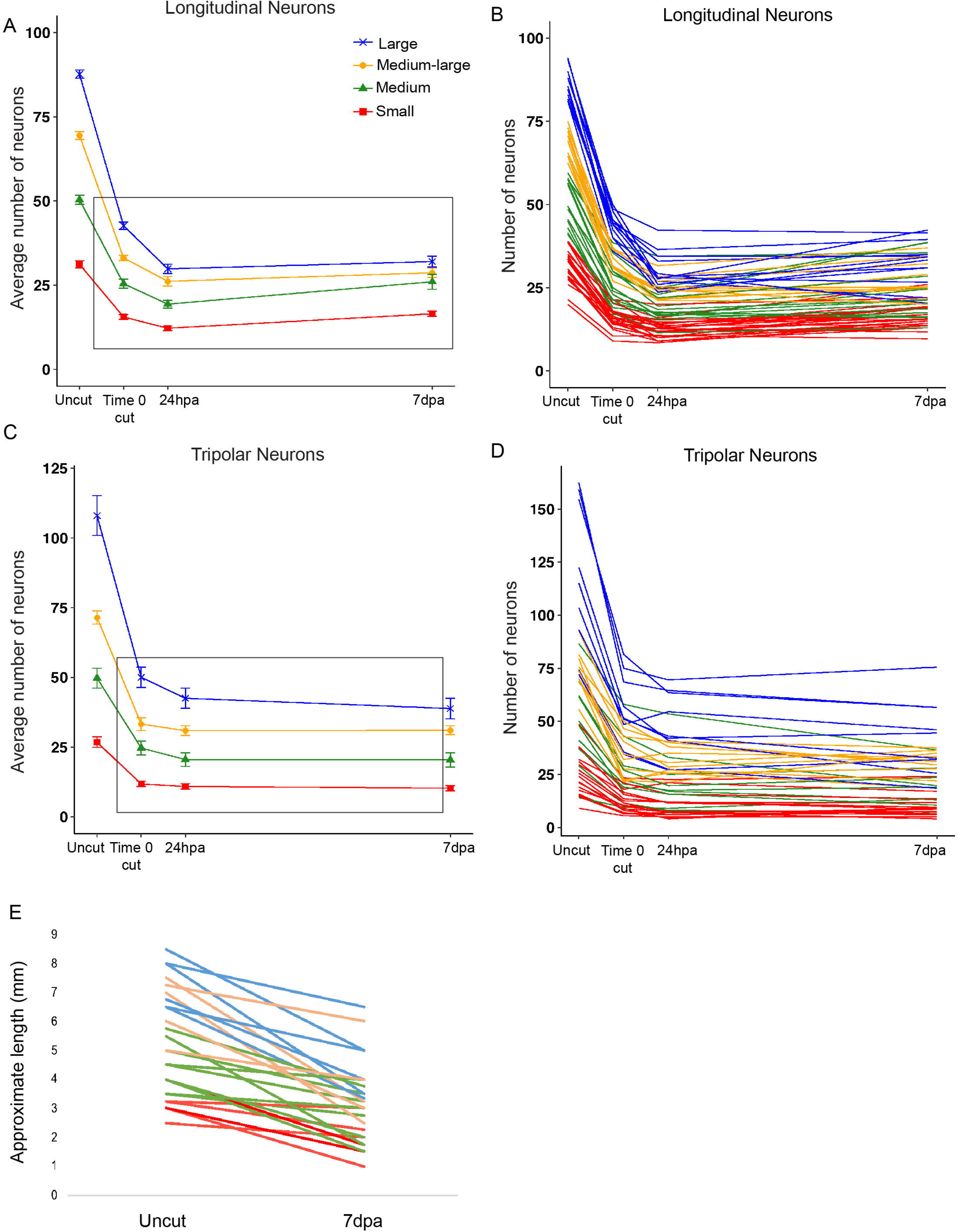
Quantification of neuronal numbers during oral regeneration. (A and C) Quantification of mean longitudinal and tripolar neurons before amputation (uncut), immediately following bisection at the oral-aboral axis midline (time 0 cut), 24 hours post amputation (hpa), and at completion of regeneration 7days post amputation (dpa). Black rectangles show time points included in analyses as seen in Figure 4F & H. (B and D) Population data showing the time course for the average number of longitudinal neurons and tripolar neurons per 2-4 radial segments in individual regenerating animals bisected in half (n = 83 animals). Neuronal numbers did not return to levels present in the uncut animal. Data represented in B and D are the same dataset grouped by starting size and averaged in A and C and Figure 4F and H. (E) Population data showing lengths for individual animals before they were amputated (uncut) and 7dpa. Regenerated animals never re-grew back to their original length (*t*_30_ = 8.65, *p* < 0.001; n = 31; see Figure 4B for percent change by size). Small animals are represented in red, medium in green, medium-large in orange, and large in blue. Data points and bars represent means ± SEM.

**Supplemental Figure 5:**
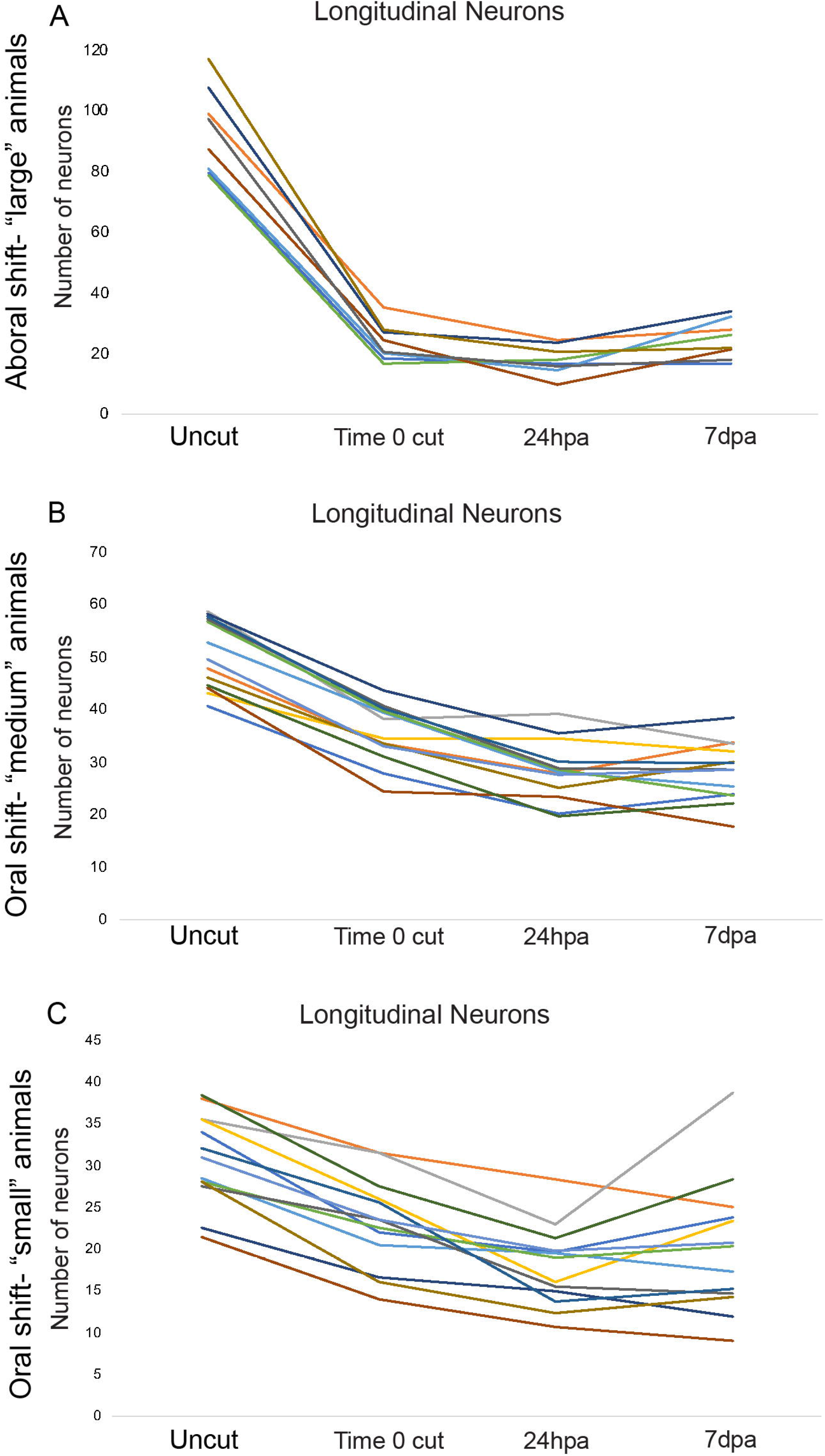
Individual longitudinal neuron data from the shifted amputation site experiments. (A) Regenerates from an aborally shifted amputation site in large animals increased their longitudinal neurons back to similar numbers present at the time of bisection (n=10) (B) Regenerates from orally shifted amputation sites in medium animals did not increase their longitudinal neurons (n=13). (C) Regenerates from orally shifted amputation sites in small animals did not increase their longitudinal neurons (n=13). Each line represents an individual animal. These data were used to determine the average responses shown in Figure 5. See Supplemental Table 5 and Figure 5 for statistical analyses.

**Supplemental Figure 6:**
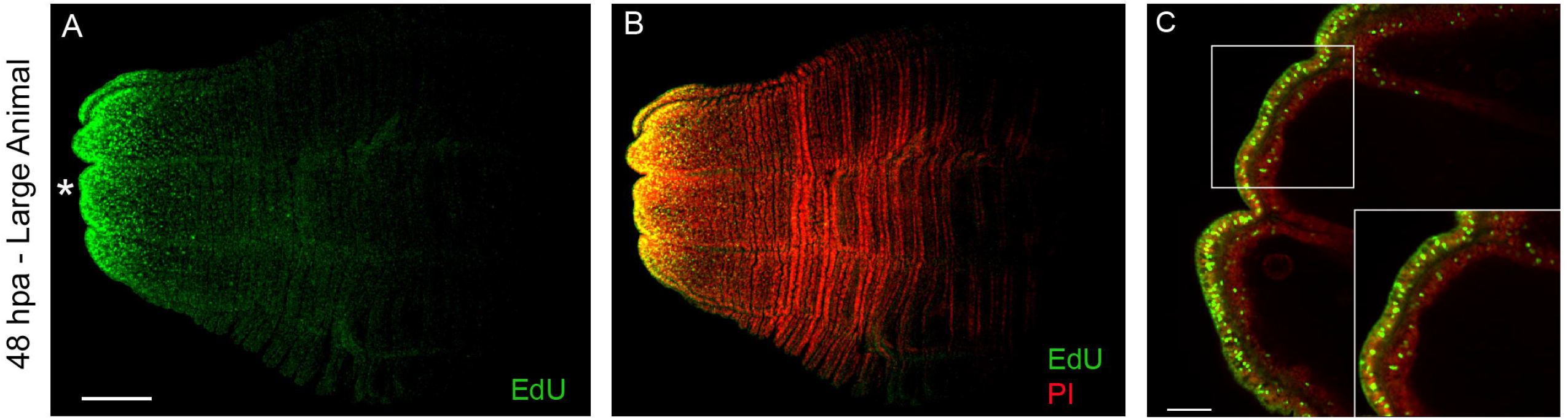
Detection of proliferating cells in “large” *Nematostella* during oral regeneration. (A) EdU (green) is detected at 48hpa in the oral-most region of large regenerating animals bisected in half. (B) EdU overlaid with propidium iodide (PI, red) stain in the same animal as A, with yellow color showing co-expression. (C) Higher magnification image of the same animal, showing EdU detection in both the ectoderm and endoderm, suggesting both tissue layers proliferate during oral regeneration (n = 3 large animals). Asterisks indicate oral opening. Scale bars for A and B = 200μm; C = 50 μm.

