## Supplemental Tables 1-4 for "Dynamics and variability of neuronal subtype responses during growth, degrowth, and regeneration"

Supplemental Table 1: Average number of tentacular neurons per tentacle in different sized animals.

| Size category | N (12 tentacles/animal) | Mean number of neurons/tentacle* | STDEV | SEM |
|---|---|---|---|---|
| Small | 3 animals | 45.22 | 25.17 | 4.20 |
| Medium | 3 animals | 71.94 | 30.57 | 5.10 |
| Medium-large | 3 animals | 84.92 | 57.13 | 9.52 |
| Large | 3 animals | 88.88 | 44.92 | 7.49 |

*Averages derived from data in supplementary table 2.

Supplemental Table 2: Number of tentacular neurons in each tentacle of animals of different sizes.

| | Size category | T1 | T2 | T3 | T4 | T5 | T6 | T7 | T8 | T9 | T10 | T11 | T12 | Min | Max |
|---|---|---|---|---|---|---|---|---|---|---|---|---|---|---|---|
| 1 | Small | 59 | 40 | 20 | 39 | 12 | 84 | 72 | 96 | 31 | 63 | 40 | 101 | 20 | 101 |
| 2 | Small | 33 | 47 | 46 | 19 | 12 | 53 | 57 | 39 | 37 | 82 | 41 | 9 | 9 | 82 |
| 3 | Small | 13 | 9 | 44 | 83 | 21 | 75 | 60 | 55 | 19 | 38 | 54 | 25 | 9 | 83 |
| 4 | Medium | 21 | 66 | 47 | 132 | 43 | 81 | 112 | 90 | 25 | 78 | 93 | 34 | 21 | 132 |
| 5 | Medium | 92 | 87 | 64 | 156 | 65 | 62 | 52 | 71 | 88 | 48 | 70 | 81 | 48 | 156 |
| 6 | Medium | 86 | 35 | 91 | 107 | 72 | 39 | 56 | 123 | 81 | 64 | 28 | 50 | 28 | 123 |
| 7 | Medium-large | 34 | 20 | 19 | 49 | 10 | 58 | 40 | 5 | 53 | 42 | 25 | 89 | 5 | 89 |
| 8 | Medium-large | 131 | 111 | 71 | 51 | 189 | 126 | 99 | 201 | 211 | 142 | 168 | 101 | 51 | 211 |
| 9 | Medium-large | 32 | 87 | 64 | 107 | 28 | 149 | 186 | 104 | 60 | 48 | 77 | 70 | 32 | 186 |
| 10 | Large | 167 | 182 | 84 | 53 | 99 | 182 | 121 | 101 | 94 | 145 | 171 | 158 | 53 | 182 |
| 11 | Large | 66 | 42 | 31 | 17 | 70 | 59 | 18 | 89 | 60 | 67 | 71 | 57 | 17 | 89 |
| 12 | Large | 129 | 51 | 102 | 104 | 50 | 89 | 99 | 51 | 34 | 102 | 96 | 89 | 34 | 129 |

T= tentacle. All animals that were counted had 12 total tentacles.

Supplemental Table 3: Number of longitudinal neurons counted in each longitudinal track.

| | Neurons counted in each of eight longitudinal tracks | | | | | | | | Mean | ± SE |
|---|---|---|---|---|---|---|---|---|---|---|
| Individual 1 | 12 | 10 | 10 | 12 | 10 | 9 | 14 | 14 | 11.38 | 0.68 |
| Individual 2 | 18 | 18 | 24 | 18 | 20 | 23 | 16 | 17 | 19.25 | 1.01 |
| Individual 3 | 41 | 39 | 40 | 35 | 38 | 38 | | | 38.5 | 0.85 |
| Individual 4 | 39 | 41 | 43 | 43 | 40 | 43 | 44 | 44 | 42.13 | 0.67 |
| Individual 5 | 47 | 46 | 49 | 45 | 47 | 46 | 44 | 42 | 45.75 | 0.71 |
| Individual 6 | 50 | 57 | 51 | 47 | 51 | 48 | 43 | 47 | 49.25 | 1.45 |
| Individual 7 | 55 | 54 | 51 | 54 | 53 | | | | 53.4 | 0.68 |
| Individual 8 | 70 | 74 | 74 | 61 | 69 | 69 | 64 | 59 | 67.5 | 1.99 |
| Individual 9 | 63 | 64 | 63 | 69 | 61 | 58 | 69 | 54 | 62.63 | 1.80 |
| Individual 10 | 69 | 67 | 75 | 74 | 67 | 65 | 64 | 67 | 68.5 | 1.41 |

Blank cells show instances where longitudinal neuron counts were not done.

Supplemental Table 4: Number of tripolar neurons counted in each radial segment.

| | Neurons counted in each of eight radial segments | | | | | | | | Mean | ± SE |
|---|---|---|---|---|---|---|---|---|---|---|
| Individual 1 | 9 | 10 | 8 | 10 | 13 | 5 | 12 | 14 | 10.13 | 1.03 |
| Individual 2 | 16 | 14 | 22 | 16 | 21 | 18 | 17 | 15 | 17.38 | 1 |
| Individual 3 | 43 | 40 | 47 | 40 | | | | | 42.5 | 1.66 |
| Individual 4 | 44 | 48 | 40 | 44 | 47 | 47 | | | 45 | 1.21 |
| Individual 5 | 65 | 50 | 70 | 57 | 45 | | | | 57.4 | 4.61 |
| Individual 6 | 52 | 55 | 51 | 57 | 51 | 48 | 43 | 47 | 50.5 | 1.58 |
| Individual 7 | 54 | 55 | 60 | 55 | 51 | | | | 55 | 1.45 |
| Individual 8 | 95 | 75 | 95 | 96 | 85 | 96 | 99 | 100 | 92.63 | 2.98 |
| Individual 9 | 87 | 86 | 85 | 75 | 76 | 75 | | | 80.67 | 2.4 |
| Individual 10 | 83 | 92 | 70 | 78 | 77 | 79 | 75 | | 79.14 | 2.61 |

Blank cells show instances where tripolar neuron counts were not done.
