## Supplemental Table 5 for "Dynamics and variability of neuronal subtype responses during growth, degrowth, and regeneration"

### Supplemental table 5: Full ANOVA model results

#### Figure 2: The *Nematostella* nerve net scales with changes in size.

Mixed ANOVA analyses were performed for the data presented in Figure 1C-E. The main effect of the repeated measure (observation time), between-subject factor (feeding regime), and the interaction effect (time x feeding) are reported for animal length (C), longitudinal neurons (D) and tripolar neurons (E). Main effects were interpreted within the context of any significant interaction effects. Bonferroni post-hoc testing was used to determine pairwise differences. S-F, starved then fed; F-S, fed then starved.

| **C) Length** | | | | | |
|---|---|---|---|---|---|
| Factors | | *Df* | *F* | *P* | $\eta_p^2$ |
| • Observation time | | 2,34 | 6.63 | **0.004** | 0.28 |
| • Feeding regime | | 1,17 | 1.02 | 0.326 | 0.06 |
| • Time x Feeding | | 2,34 | 25.75 | **< 0.001** | 0.602 |
| *Pairwise comparisons:* | | | | *Mean difference* | *P* |
| *(Feeding by Time)* | | | | | |
| Week 1: | S-F vs. F-S | | | 0.22 | 0.280 |
| Feeding regime switch: | S-F vs. F-S | | | - 0.62 | **0.003** |
| Week 14: | S-F vs. F-S | | | 0.91 | **0.002** |
| *(Time by Feeding)* | | | | | |
| S-F: | Week 1 vs. Feed switch | | | 0.32 | 0.098 |
| | Week 1 vs. Week 14 | | | - 0.72 | **0.002** |
| | Feed switch vs. Week 14 | | | - 1.04 | **< 0.001** |
| F-S: | Week 1 vs. Feed switch | | | - 0.52 | **0.008** |
| | Week 1 vs. Week 14 | | | - 0.03 | 1.00 |
| | Feed switch vs. Week 14 | | | 0.49 | **0.008** |
| **D) Number of longitudinal neurons** | | | | | |
| Factors | | *Df* | *F* | *P* | $\eta_p^2$ |
| • Observation time | | 2,36 | 9.91 | **< 0.001** | 0.36 |
| • Feeding regime | | 1,18 | 0.52 | 0.82 | 0.003 |
| • Time x Feeding | | 2,36 | 77.35 | **< 0.001** | 0.81 |
| *Pairwise comparisons:* | | | | *Mean difference* | *P* |
| *(Feeding by Time)* | | | | | |
| Week 1: | S-F vs. F-S | | | 18.3 | 0.43 |
| Feeding switch: | S-F vs. F-S | | | - 129.6 | **0.001** |
| Week 14: | S-F vs. F-S | | | 127.7 | **< 0.001** |
| *(Time by Feeding)* | | | | | |
| Starved then fed: | Week 1 vs. Feed switch | | | 44.6 | **0.007** |
| | Week 1 vs. Week 14 | | | - 100.3 | **< 0.001** |
| | Feed switch vs. Week 14 | | | - 144.8 | **< 0.001** |
| Fed then starved | Week 1 vs. Feed switch | | | - 103.3 | **< 0.001** |
| | Week 1 vs. Week 14 | | | 9.1 | 1.00 |
| | Feed switch vs. Week 14 | | | 112.4 | **< 0.001** |

| **E) Number of tripolar neurons** | | | | |
|---|---|---|---|---|
| Factors | *Df* | *F* | *P* | $\eta_p^2$ |
| • Observation time | 2,36 | 20.69 | **< 0.001** | 0.54 |
| • Feeding regime | 1,8 | 2.09 | 0.19 | 0.21 |
| • Time x Feeding | 2,36 | 91.91 | **< 0.001** | 0.84 |

| *Pairwise comparisons:* | | *Mean difference* | *P* |
|---|---|---|---|
| *(Feeding by Time)* | | | |
| Week 1: | S-F vs. F-S | 13.4 | 0.61 |
| Feeding regime switch: | S-F vs. F-S | - 115.1 | **< 0.001** |
| Week 14: | S-F vs. F-S | 184.6 | **< 0.001** |
| *(Time by Feeding)* | | | |
| S-F: | Week 1 vs. Feed switch | 64.9 | **0.03** |
| | Week 1 vs. Week 14 | - 147.5 | **< 0.001** |
| | Feed switch vs. Week 14 | - 212.4 | **< 0.001** |
| F-S: | Week 1 vs. Feed switch | - 63.6 | **0.032** |
| | Week 1 vs. Week 14 | 23.7 | 0.70 |
| | Feed switch vs. Week 14 | 87.3 | **0.03** |

**<u>Figure 4: Differential regeneration of *NvLWamide-like* neurons depends on the specific neuronal subtype</u>.** Mixed ANOVA analyses were performed for the data presented in Figure 4F&H. The main effect of the repeated measure (observation time), between-subject factor (starting size category), and the interaction effect (time x size) are reported for longitudinal (F) and tripolar neurons (H). Main effects were interpreted within the context of any significant interaction effects. Bonferroni post-hoc testing was used to determine pairwise differences. Greenhouse-Geisser corrected *F* statistics are reported for longitudinal and tripolar data sets (F, H), due to a lack of sphericity. Dpa, days post amputation; hpa, hours post amputation.

| **F) Regeneration of longitudinal neurons** | | | | |
|---|---|---|---|---|
| Factors | *Df* | *F* | *P* | $\eta_p^2$ |
| • Observation time | 1.7,134.4 | 83.95 | **< 0.001** | 0.52 |
| • Starting size | 3,79 | 79.11 | **< 0.001** | 0.75 |
| • Time x Size | 5.1,134.4 | 13.46 | **< 0.001** | 0.34 |

| *Pairwise comparisons:* | | *Mean difference* | *P* |
|---|---|---|---|
| *(Time by Size)* | | | |
| Small: | Time 0 cut vs. 24 hpa | 3.3 | **< 0.001** |
| | Time 0 cut vs. 7 dpa | - 1.0 | 0.99 |
| | 24 hpa vs. 7 dpa | - 4.3 | **< 0.001** |
| Medium: | Time 0 cut vs. 24 hpa | 6.1 | **< 0.001** |
| | Time 0 cut vs. 7 dpa | -0.6 | 1.00 |
| | 24 hpa vs. 7 dpa | - 6.6 | **< 0.001** |
| Medium-large: | Time 0 cut vs. 24 hpa | 6.9 | **< 0.001** |
| | Time 0 cut vs. 7 dpa | 4.4 | **0.003** |
| | 24 hpa vs. 7 dpa | - 2.5 | 0.25 |
| Large: | Time 0 cut vs. 24 hpa | 12.8 | **< 0.001** |
| | Time 0 cut vs. 7 dpa | 10.6 | **< 0.001** |
| | 24 hpa vs. 7 dpa | - 2.1 | 0.34 |
| *(Size by Time)* | | | |
| Time 0 cut: | Small vs. Medium | - 9.9 | **< 0.001** |
| | Small vs. Medium-large | - 17.5 | **< 0.001** |
| | Small vs. Large | - 27.1 | **< 0.001** |
| | Medium vs. Medium-large | - 7.7 | **< 0.001** |
| | Medium vs. Large | - 17.2 | **< 0.001** |
| | Medium-large vs. Large | - 9.5 | **< 0.001** |
| 24 hpa: | Small vs. Medium | - 7.2 | **< 0.001** |
| | Small vs. Medium-large | -14.0 | **< 0.001** |
| | Small vs. Large | - 17.6 | **< 0.001** |
| | Medium vs. Medium-large | - 6.8 | **< 0.001** |
| | Medium vs. Large | - 10.5 | **< 0.001** |
| | Medium-large vs. Large | - 3.7 | 0.19 |
| 7 dpa: | Small vs. Medium | - 9.5 | **< 0.001** |
| | Small vs. Medium-large | - 12.2 | **< 0.001** |
| | Small vs. Large | - 15.5 | **< 0.001** |
| | Medium vs. Medium-large | - 2.7 | 1.00 |
| | Medium vs. Large | - 6.0 | **0.05** |
| | Medium-large vs. Large | - 3.3 | 0.96 |

**H) Regeneration of tripolar neurons**

| Factors | *Df* | *F* | *P* | $\eta_p^2$ |
|---|---|---|---|---|
| • Observation time | 1.5,116.2 | 22.75 | **< 0.001** | 0.23 |
| • Starting size | 3,77 | 54.18 | **< 0.001** | 0.68 |
| • Time x Size | 4.5,116.2 | 3.93 | **0.003** | 0.13 |

| *Pairwise comparisons:* | | *Mean difference* | *P* |
|---|---|---|---|
| *(Time by Size)* | | | |
| Small: | Time 0 cut vs. 24 hpa | 0.9 | 1.00 |
| | Time 0 cut vs. 7 dpa | 1.2 | 1.00 |
| | 24 hpa vs. 7 dpa | 0.4 | 1.00 |
| Medium: | Time 0 cut vs. 24 hpa | 4.4 | **0.001** |
| | Time 0 cut vs. 7 dpa | 4.2 | 0.08 |
| | 24 hpa vs. 7 dpa | - 0.2 | 1.00 |
| Medium-large: | Time 0 cut vs. 24 hpa | 3.8 | **0.013** |
| | Time 0 cut vs. 7 dpa | 3.0 | 0.46 |
| | 24 hpa vs. 7 dpa | - 0.8 | 1.00 |
| Large: | Time 0 cut vs. 24 hpa | 6.8 | **< 0.001** |
| | Time 0 cut vs. 7 dpa | 10.2 | **< 0.001** |
| | 24 hpa vs. 7 dpa | 3.4 | 0.055 |
| *(Size by Time)* | | | |
| Time 0 cut: | Small vs. Medium | - 13.6 | **< 0.001** |
| | Small vs. Medium-large | - 23.9 | **< 0.001** |
| | Small vs. Large | - 38.3 | **< 0.001** |
| | Medium vs. Medium-large | - 10.3 | **0.03** |
| | Medium vs. Large | - 24.7 | **< 0.001** |
| | Medium-large vs. Large | - 14.4 | **0.001** |
| 24 hpa: | Small vs. Medium | - 10.1 | **0.003** |
| | Small vs. Medium-large | -21.1 | **< 0.001** |
| | Small vs. Large | - 32.4 | **< 0.001** |
| | Medium vs. Medium-large | - 11.0 | **0.005** |
| | Medium vs. Large | - 22.3 | **< 0.001** |
| | Medium-large vs. Large | - 11.3 | **0.004** |
| 7 dpa: | Small vs. Medium | - 10.6 | **0.003** |
| | Small vs. Medium-large | - 22.2 | **< 0.001** |
| | Small vs. Large | - 29.3 | **< 0.001** |
| | Medium vs. Medium-large | - 11.6 | **0.005** |
| | Medium vs. Large | - 18.7 | **< 0.001** |
| | Medium-large vs. Large | - 7.1 | 0.22 |

**Figure 5: Regeneration of longitudinal neurons depends on the size of the regenerating fragment.**
Repeated measure ANOVA analyses were performed for the data presented in Figure 5C, E, G. Observation time served as the repeated measure for large animals with an aboral shift in cut site (C), medium animals with an oral shift in cut site (E), and small animals with an oral shift in cut site (G). Bonferroni post-hoc tests were used to evaluate pairwise differences when there was a significant main effect of observation time on the number of neurons observed. Dpa, days post amputation; hpa, hours post amputation.

| **C) Regeneration of large animals with aboral shift in cut site** | | | | |
|---|---|---|---|---|
| Factors | *Df* | *F* | *P* | $\eta_p^2$ |
| • Observation time | 2,14 | 5.58 | **0.017** | 0.44 |
| *Pairwise comparisons:* | | | *Mean difference* | *P* |
| Time 0 cut vs. 24 hpa | | | 5.8 | **0.04** |
| Time 0 cut vs. 7 dpa | | | - 1.0 | 1.00 |
| 24 hpa vs. 7 dpa | | | - 6.9 | **0.046** |
| **E) Regeneration of medium animals with oral shift in cut site** | | | | |
| Factors | *Df* | *F* | *P* | $\eta_p^2$ |
| • Observation time | 2,24 | 21.67 | **< 0.001** | 0.64 |
| *Pairwise comparisons:* | | | *Mean difference* | *P* |
| Time 0 cut vs. 24 hpa | | | 7.0 | **< 0.001** |
| Time 0 cut vs. 7 dpa | | | 7.1 | **0.001** |
| 24 hpa vs. 7 dpa | | | 0.05 | 1.00 |
| **G) Regeneration of small animals with oral shift in cut site** | | | | |
| Factors | *Df* | *F* | *P* | $\eta_p^2$ |
| • Observation time | 2,24 | 8.56 | **0.002** | 0.42 |
| *Pairwise comparisons:* | | | *Mean difference* | *P* |
| Time 0 cut vs. 24 hpa | | | 5.1 | **< 0.001** |
| Time 0 cut vs. 7 dpa | | | 2.9 | 0.12 |
| 24 hpa vs. 7 dpa | | | -2.2 | 0.47 |

**Supplemental Figure 4: Length measurements and neural quantifications in *Nematostella*.**
A one-way ANOVA was performed for the data presented in Supplemental Figure 4F. (F) Percent length regenerated by 7dpa was evaluated using shifted cut site location in small, medium and large animals as a factor. Bonferroni post-hoc testing was used to evaluate pairwise differences.

| **F) Percent length regenerated at 7dpa based on cutsite in different sized animals** | | | | |
|---|---|---|---|---|
| Factors | *Df* | *F* | *P* | $\eta_p^2$ |
| • Cut site | 2,28 | 10.92 | **< 0.001** | 0.44 |
| *Pairwise comparisons:* | | | *Mean difference* | *P* |
| Small animal oral cut vs. Medium animal oral cut | | | - 0.03 | 1.00 |
| Small animal oral cut vs. Large animal aboral cut | | | 0.25 | **0.001** |
| Medium animal oral cut vs. Large animal aboral cut | | | 0.28 | **< 0.001** |
