## Supplemental Movie Legends for "Dynamics and variability of neuronal subtype responses during growth, degrowth, and regeneration"

**Supplemental movie 1: Response to poking assay prior to prolonged starvation.** Poking assays were performed prior to starvation (time 0). This *Nematostella* individual responded to the touch stimulus by retracting its tentacles. Video duration: 00:07; 3 frames/second.

**Supplemental movie 2: Response to poking assay is maintained after prolonged starvation.** Poking assays were performed following prolonged starvation. This *Nematostella* individual was starved for 7 weeks and responded to the touch stimulus by retracting its tentacles. This video is of the same individual as Supplemental movie 1. Video duration: 00:06; 3 frames/second.

**Supplemental movie 3: Regenerated animals can capture and ingest prey.** Feeding assays were performed by providing live artemia to individuals following regeneration. This *Nematostella* individual was fed 7 days post amputation and captured and ingested multiple artemia. Video duration: 00:08; 100 frames/second.

**Supplemental movie 4: Regenerated animals can digest ingested prey.** Feeding assays were performed by providing live artemia to individuals following regeneration. This *Nematostella* individual was fed 7 days post amputation and two artemia can be observed digesting inside the body cavity. This video is of the same individual as Supplemental movie 3 but shows the animal 90 minutes later. Video duration: 00:06; 100 frames/second.

**Supplemental movie 5: Regenerated animals perform peristaltic waves.** Animals were observed for their ability to propagate peristaltic waves down their body columns following regeneration. This *Nematostella* individual was observed 7 days post amputation and could perform peristaltic wave behaviors. Video duration: 00:04; 100 frames/second.

**Supplemental movie 6: Response to poking assay prior to bisection in half.** Poking assays were performed prior to amputation (time 0 uncut). This *Nematostella* individual responded to the touch stimulus by retracting its tentacles. Video duration: 00:08; 3 frames/second.

**Supplemental movie 7: Response to poking assay is maintained in regenerating animals.** Poking assays were performed following regeneration (7dpa). This *Nematostella* individual responded to the touch stimulus by retracting its tentacles. This video is of the same individual as Supplemental movie 1. Video duration: 00:07; 3 frames/second.
